# Meta-analysis of spatial genetic patterns among European saproxylic beetles - the influence of history and contemporary forest quality

**DOI:** 10.1101/2024.07.24.604895

**Authors:** Rama Sarvani Krovi, Nermeen Amer, Maria Oczkowicz, Łukasz Kajtoch

## Abstract

The phylogeography of forest-dwelling species in Europe is well understood, although our knowledge regarding the genetics of saproxylic beetles remains insufficient. This knowledge gap extends to understanding the influence of both quaternary history and contemporary forest dynamics on population genetics. To fill this gap, we conducted a systematic review and meta-analysis of recent literature concerning saproxylic beetle taxa with available genetic data. We include both threatened and common species in our study which enabled us to generalize our findings to the whole saproxylic community. Results suggest a latitudinal decrease in diversity in most species, likely influenced by Pleistocene glaciation and subsequent population expansions from southern refugia. Additionally, we observed an east-west gradient in diversity, with threatened species exhibiting higher diversity towards the east. This may reflect historical forest dynamics and anthropogenic pressures, such as heavy wood logging in Western Europe. Similarly, we found a pattern along altitude, with populations in higher elevation forests, which are often more natural, exhibiting higher diversity. Furthermore, we identified distinct phylogenetic units or genetic clusters in southern Europe reflecting the distribution of glacial refugia. For some taxa, distinct units were also reported in eastern Europe where populations spread from Asian refugia. Central Europe showed a high number of phylogenetic units, although unique (private) clades or clusters were absent. Most likely it is an effect of the presence of beetles that originated from various refugia belonging to different phylogenetic units.

This study brings insights into general phylogeographic patterns, which have previously been examined only for single representatives of saproxylic beetles. It should also help in the proper planning of conservation and management efforts of wood-dwelling beetles.

## 1. Introduction

Forest-dwelling organisms are one of the most diverse groups of terrestrials in the world as many areas are, or at least used to be, covered by trees (Erwin, 1997, Lieutier et al, 2004). Trees cover huge areas from the tropics to the boreal zone, except for areas with extreme temperatures and water deficiency (subtropical, arctic, and mountain deserts). The distribution and diversity of forests were shaped by natural conditions like climate, water supplies, and soils. The latest history of trees was highly associated with glacial cycles, which forced forests to retreat to tropic zones or refugial areas in higher latitudes (Taberlet et al., 1998). In Europe, such refugia were mostly situated in the Mediterranean Basin (Hewitt et al, 1999; Kerdelhué et al., 2002), although numerous local refugia were also discovered in Western and Central Europe (Stewart et al., 2001). Some of the trees (and associated species) found suitable areas for ice-age survival in the eastern part of Europe or Asia. Consequently, the phylogeography of European trees and arboreal taxa is highly diverse, and apart from major patterns (following paradigms of other temperate species), there are many exceptions (Taberlet et al.,1998; Stewart et al., 2010).

Later, during the Holocene, trees expanded and formed the dominant ecosystems in Europe: Mediterranean, temperate, and boreal forests (Marquer et al., 2017). Various tree species, either coniferous or deciduous, formed particular forest types, rarely being monotypic. An important element of forests is the wood of dead trees, which is particularly noticeable in temperate and boreal zones, due to the long time needed for wood to decay (in contrast to tropics) (Harmon et al., 1986) A large number, volume and various stages of wood of dead trees enable the formation of very rich assemblages of saproxylic taxa among which many evolved to utilize the common substrate – wood (Hjältén et al., 2012). Many other taxa living in the wood of dead trees selected other food sources like widespread fungi or eating live or dead saprophagous invertebrates. This resulted in very rich communities, with the most diverse like fungi, lichens, bryophytes, and many animals (both invertebrates and vertebrates) (Stokland et al., 2012).

This ecosystem was next modified by humans, who found wood a very important resource (e.g. as fuel, or constructions), or removed trees to transform land for other uses (agriculture, settlement etc.) (Taylor et al., 2009). Changes in woodland cover and forest quality caused by human activity since the antiquity to industrial times forced many tree-living species to change their ranges, and abundance or cause extinction (Ulrich, 1995; Thiollay, 2006). On the other hand, the spread of planted trees enables the expansion of other organisms, particularly those that are considered pests in forestry (Freer-Smith et al., 2019). The decline of populations of rare species and the outbreak of common ones are now accelerated by climate changes, which have a serious impact also on forests all over the world (Bauhus et al, 2017).

Among species living in forests, the crucial group constitutes taxa utilizing wood as a microhabitat for living and/or foraging (Hoffman et al, 2000). Many of these species depend on the amount and quality of wood from dead trees (colloquially “deadwood”), which is a natural element in forest ecosystems (Siitonen, 2001). It is a common substrate in natural forests, as trees are dying and naturally their remains are left in the ecosystem for decades, making space for numerous species living in or on the wood of dead trees. This is rare in the case of managed (commercial) forests, where wood production is a major activity, so most of the dead trees are removed before they can be utilized by these species. Consequently, many organisms that depend on the so-called deadwood, cannot survive in planted and logged forests, so their populations in such forests are scattered and small (Freedman et al, 2011). Again, this is not a rule, as some organisms adapt to such heavily transformed habitats and start to be abundant, which sometimes leads to their outbreaks having an impact on the economic value of forests (Klapwijk et al. 2016).

Beetles (Coleoptera) are likely the best-known group of saproxylic organisms at least in temperate and boreal forests in the Northern Hemisphere (Gimmel et al., 2018). Generally, beetles are the most diverse group of animals (insects) in the world and many of them are associated with forests, with numerous taxa living in various forms of dying and dead trees (Ślipiński et al., 2011; Gimmel & Ferro., 2018). They play an important role in wood decomposition and in the cycling of nutrients, carbon, and microelements in forest ecosystems (Jomura et al., 2020). The widest knowledge is for taxa living in Europe, North America as well as in eastern Asia, although it is highly unequal (Kajtoch et al., 2022). So-called “pests’’ are objects of numerous studies all over the world, with the greatest focus on taxa having economic value for coniferous forests in the Holarctic (De Groot et al. 2019). On the other hand, attention to rare and threatened taxa is also increasing for European taxa, especially thanks to the development of red lists of saproxylic beetles (Carpaneto et al. 2015). The overall knowledge is also unbalanced as most available information is related to biogeography (distribution), and ecology (habitat/microhabitat requirements, trophism). Data about genetic polymorphism of saproxylic beetles is even weaker as not much research was done on members of this group (Kajtoch et al. 2022). The exceptions are “pests’’ and invasive taxa, which were relatively frequent objects of genetic studies, but again, usually on only some levels (like phylogenetic or delimitation for some groups, e.g. bark beetles), or for only some populations (e.g. mostly these from western, southern or northern Europe or western North America) (Kajtoch et al. 2022). Genetic data about the majority of common species are still unknown. Even for rare and threatened beetles not so many genetic studies are available, and usually they were focused on only some charismatic species or some populations. Kajtoch et al. (2022) provided a comprehensive review of the genetic diversity of saproxylic beetles worldwide. However, the review was too general to address numerous questions adequately, revealing more unexplored areas ("white spots"). Overall, it suggested that glacial and post-glacial range shifts primarily shaped genetic variability in saproxylic beetle populations. Additionally, certain species appeared to be linked to habitat quality and quantity which is important for forest conservation or management strategies.

Considering the above, a list of relevant studies was searched in the literature to solve the hypothesis assuming that the genetic diversity of saproxylic beetle populations decreased to the north as it is in most wood-dwelling species in Europe. Similarly, we could assume that for boreal taxa genetic polymorphism should be higher in the eastern populations (being less affected by glacials). Moreover, the polymorphism of populations on higher altitudes should be higher due to the quality of forests being less affected by timber logging in mountains.

Second, the number and distribution of genetic units (either mitochondrial clades or microsatellite clusters) were compared among species to find generalities (e.g. presence of distinct units in various regions of Europe, that could have either conservation or management value).

Finally, the genetic diversity of populations of rare taxa should have overall lower polymorphism than pest taxa, as the latter is less (or not) affected by forest management actions.

The general idea of this review is to summarize the knowledge on genetic polymorphism of saproxylic beetles being studied in Europe to give a background for effective planning of conservation (for rare and threatened taxa), or management (for taxa having economic value in wood production). This review and meta-analysis should be also valuable background for future studies on the genetic variability of saproxylic beetles.

## 2. Methods

### 2.1. Literature search

The literature search was conducted according to PRISMA (Preferred Reporting Items for Systematic Reviews and Meta-Analyses) guidelines in the WoS (Web of Science) and Scopus databases. The search was done exactly like in a review of (Kajtoch et al., 2022) but extended to recent publications (2022-2023, till 31.12.2023). The same criteria were adopted to newest papers search with use of the same keywords: *(ALL=(Phylogeny OR phylogenetic OR phylogeography OR phylogeographic OR population genetic OR conservation genetic OR landscape genetic OR population genomic OR conservation genomic OR landscape genomic) ALL=(AND beetle OR Coleoptera AND saproxylic OR xylophagic OR saproxylophagic OR cambiophagic OR cambioxylophagic OR xylophagous OR saproxylophagous OR cambiophagous OR cambioxylophagous OR deadwood))*. All relevant papers were retrieved according to the following criteria: i) concern species from Europe, ii) available data for numerous sites across the continent with georeferences, iii) available genetic polymorphism metrics (haplotype diversity (Hd) based on mitochondrial DNA markers, and/or heterozygosity (Ho) based on microsatellites). Heterozygosity (observed heterozygosity) was selected as more articles reported this metric than allelic richness (moreover allelic richness is more dependet on sample size). The same was in case of haplotype diversity being more frequently available in articles than nucleotide diversity. Due to a very few studies based on the genomic data (from next-generation sequencing) (Schebeck et al. 2018, 2019, Ellerstand et al 2022, Mykhailenko et al. (in press)), in this review, we resigned from the presentation of genetic metrics measured from single nucleotide polymorphisms.

### 2.2. Geographic assignment of populations

Initially, it was planned to analyze intraspecific variability in particular populations or their groups in Europe separately, but due to the large differences in several data (examined sites), the analysis was restricted to a species level. For some species with available mitochondrial or microsatellite data showing overall intraspecific diversity in Europe, the general picture of genetic population structure (number and distribution of clades or clusters) was presented. The same was done for the presence of identified refugia across defined regions of Europe and adjacent areas, namely: Scandinavian (Norway, Sweden, Denmark, Finland), North-Western (Ireland, Britain), Western (Benelux, lowland France, and Germany), Iberian (Portugal, Spain), Appennines (S Italy), Balkans (Balkan Peninsula), Alpine (Switzerland, Austria, Slovenia, S Germany, N Italy, SE France), Pannonian-Carpathian-Bohemian (Czech R., Slovakia, S Poland, SE Ukraine, inner Romania, Hungary), Pontic (E Romania, Moldova, S Ukraine, SW Russia), North-Eastern (N Poland, Baltic countries,), Eastern (W, N, central Russia Belarus), Caucasus (Georgia, Armenia, Azerbaijan, SE Russia), and Anatolia (Turkey).

### 2.3. Statistical analyses

All statistical analyses were implemented using R (R Core Team 2020, R Studio Team 2015, Wickham, 2016). For genetic diversity (Hd and Ho), we ran separate models for Hd and Ho with species, longitude, latitude, and altitudes and their interactions as predictors. Subsequently, separate models for each saproxylic species separately were performed with longitude, latitude, and altitude as predictors. All models were performed using a generalized linear model (glm function) and Gaussian distribution. P-values were obtained using ANOVA implemented in the car package (Fox and Weisberg 2015). Selection of the best model was made based on the corrected Akaike’s information criteria for sample size (AICc) and weights. Then, we ran a specific model based on the output of the model selection analysis. In case AICc returns “Null model”, we chose the next best model with (AIC < 2). Relationships between geographical dimensions (longitude, latitude, and altitude) with haplotype diversity or heterozygosity were assessed using linear associations (Pearson, 1895). All boxplots and scatterplots were incorporated using (ggplot2) for visualization. All maps were conducted using QGIS software (version: 3.36.3, 2019)

## 3. Results

The output for our literature search resulted in 1734 publications among which 167 included papers referring to the genetics of European saproxylic beetles but only 60 included phylogeographic or population genetic information. After selection, 31 publications were used for meta-analysis and a further 13 papers were for descriptive examination a long with sample sizes (Table S1). Data preprocessing involved removing duplicates, handling missing values, and verifying location coordinates’ accuracy along with generating altitudes from the coordinates provided in the study. (Table S2).

The goals of these studies were mostly focused on phylogeography, population genetics or conservation genetics, and evolutionary or delimitation (with barcoding) topics were included. Noticeably, no research includes environmental data (e.g. forest structure, continuity, or quality).

### 3.1. General overview

Twenty species of saproxylic beetles were considered for this review. Information on genetic diversity data was available for 11 species (nine with mitochondrial data and five with microsatellites). Moreover, for all 20 taxa general phylogeographic and/or population genetic information were available, which were used for descriptive elaboration of beetle history and distribution of evolutionary units. Among selected beetles are both threatened species (*Osmoderma* spp., *Rosalia alpina, Morimus funereus, Cucujus cinnaberinus, Pytho kolwensis*), and common taxa, either of economic value (e.g. *Ips typographus, Ips sexdentatus, Pityogenes chalcographus, Tomicus destruens, Tomicus piniperda*), or being neutral in forestry (e.g. *Cetonia aurata, Oxythyrea spp., Bolitophagus reticulatus, Lucanus cervus, Rhagium inquisitor, Monochamus sartor, Morimus asper, Anastrangalia spp., Pytho depressus, Pytho abieticola*) (Table 1).

**Table 1.** Species considered for this review and the conservation status. Studies were taken between the years 2000 and 2023.

| Family | Species | Reference | Genetic marker examined | Data used | conservation status & protection | considered as pest |
| --- | --- | --- | --- | --- | --- | --- |
| Scarabeidae | *Cetonia aurata* | Ahrens et al. (2013) | mtDNA | general phylogeography | not threatened - not protected | no |
| Scarabeidae | *Osmoderma spp.* | Audisio et al. (2009) | mtDNA | general phylogeography | vulnerable - protected | no |
| Scarabeidae | *Oxythyrea spp.* | Vondracek et al. (2023) | mtDNA | general phylogeography | not threatened - not protected | no |
| Tenebrionidae | *Bolitophagus reticulatus* | Eberle et al. (2021) | mtDNA, microsatellite | general phylogeography & genetic diversity meta-analysis | not threatened - not protected | no |
| Lucainidae | *Lucanus cervus* | Solano et al. (2016) | mtDNA, microsatellite | general phylogeography & genetic diversity meta-analysis | near threatened - protected | no |
| Cerambycidae | *Rhagium inquisitor* | Cakmak et al. (2020) | mtDNA | general phylogeography | not threatened - not protected | sometimes |
| Cerambycidae | *Rosalia alpina* | Molfini et al. (2018) | mtDNA, microsatellite | general phylogeography & genetic diversity meta-analysis | endangered - protected | no |
| Cerambycidae | *Monochamus sartor* | Plewa et al. (2018) | mtDNA, microsatellite | general phylogeography & genetic diversity meta-analysis | not threatened - not protected | no |
| Cerambycidae | *Morimus asper* | Gojkovic et al. (2022) | mtDNA | general phylogeography | not threatened - not protected | no |
| Cerambycidae | *Morimus funereus* | Solano et al. (2013) | mtDNA | general phylogeography | not threatened - not protected | no |
| Cerambycidae | *Anastrangalia spp.* | Zamoroka et al. (2019) | mtDNA | general phylogeography | not threatened - not protected | no |
| Cucujidae | *Cucujus cinnaberinus* | Sikora et al. (2023) | mtDNA,<br>microsatellite | general phylogeography<br>& genetic diversity meta-analysis | vulnerable -<br>protected | no |
| Pythidae | *Pytho kolwensis* | Painter et al. (2007) | mtDNA | general phylogeography | vulnerable -<br>protected | no |
| Pythidae | *Pytho depressus* | Painter et al. (2007) | mtDNA | general phylogeography | not threatened - not<br>protected | no |
| Pythidae | *Pytho abieticola* | Painter et al. (2007) | mtDNA | general phylogeography | not threatened - not<br>protected | no |
| Curculionidae | *Ips sexdentatus* | Avtzis et al. (2019) | mtDNA | general phylogeography | not threatened - not<br>protected | yes |
| Curculionidae | *Ips typographus* | Stauffer et al. (1999) | mtDNA,<br>microsatellite | general phylogeography<br>& genetic diversity meta-analysis | not threatened - not<br>protected | yes |
| Curculionidae | *Pityogenes chalcographus* | Schebeck et al. (2018) | mtDNA | general phylogeography<br>& genetic diversity meta-analysis | not threatened - not<br>protected | yes |
| Curculionidae | *Tomicus destruens* | Kerdelhue et al. (2002) | mtDNA | general phylogeography<br>& genetic diversity meta-analysis | not threatened - not<br>protected | yes |
| Curculionidae | *Tomicus pinniperda* | Kerdelhue et al. (2002) | mtDNA | general phylogeography<br>& genetic diversity meta-analysis | not threatened - not<br>protected | yes |

### 3.2. Genetic diversity and sampling sites distribution

Haplotype diversity was available for 12 beetle species. Although, for three species, the number of independent sites was too low for any analyses (namely: *Monochamus galloprovincialis* (Cerambycidae; Vondracek et al, 2018), and *Osmoderma barnabita* (Scarabaeidae; Melosik et al, 2020). For *Bolitophagus reticulatus* (Eberle et al., 2021) information about haplotype diversities were provided for groups of populations from a given country, therefore it was not possible to use these data in most analyses. Two pairs of sibling species: *Tomicus pinniperda* and *T. destruens* (Curculionidae; Kerdelhué et al., 2002;), were formerly considered as one, and belong to the species complex. The final list of examined species for haplotype diversities includes eight taxa, that are: *Lucanus cervus* (Lucanidae; Solano et al, 2015), *Rosalia alpina* (Cerambycidae; Drag et al, 2018), *Cucujus cinnaberinus* (Cucujidae; Sikora et al., 2022), *Protaetia cuprea* (Scarabaeidae; Vondráček et al., 2018), *Monochamus sartor* (Cerambycidae; Plewa et al, 2018), *Ips typographus* (Curculionidae; Stauffer et al., 1999;), *Pityogenes chalcographus* (Curculionidae; Schebeck et al., 2018), *Tomicus destruens* and *T. pinniperda* (Curculionidae; Faccoli et al., 2005) (Fig. 1, Table 1, Table S3).

**Fig. 1.**
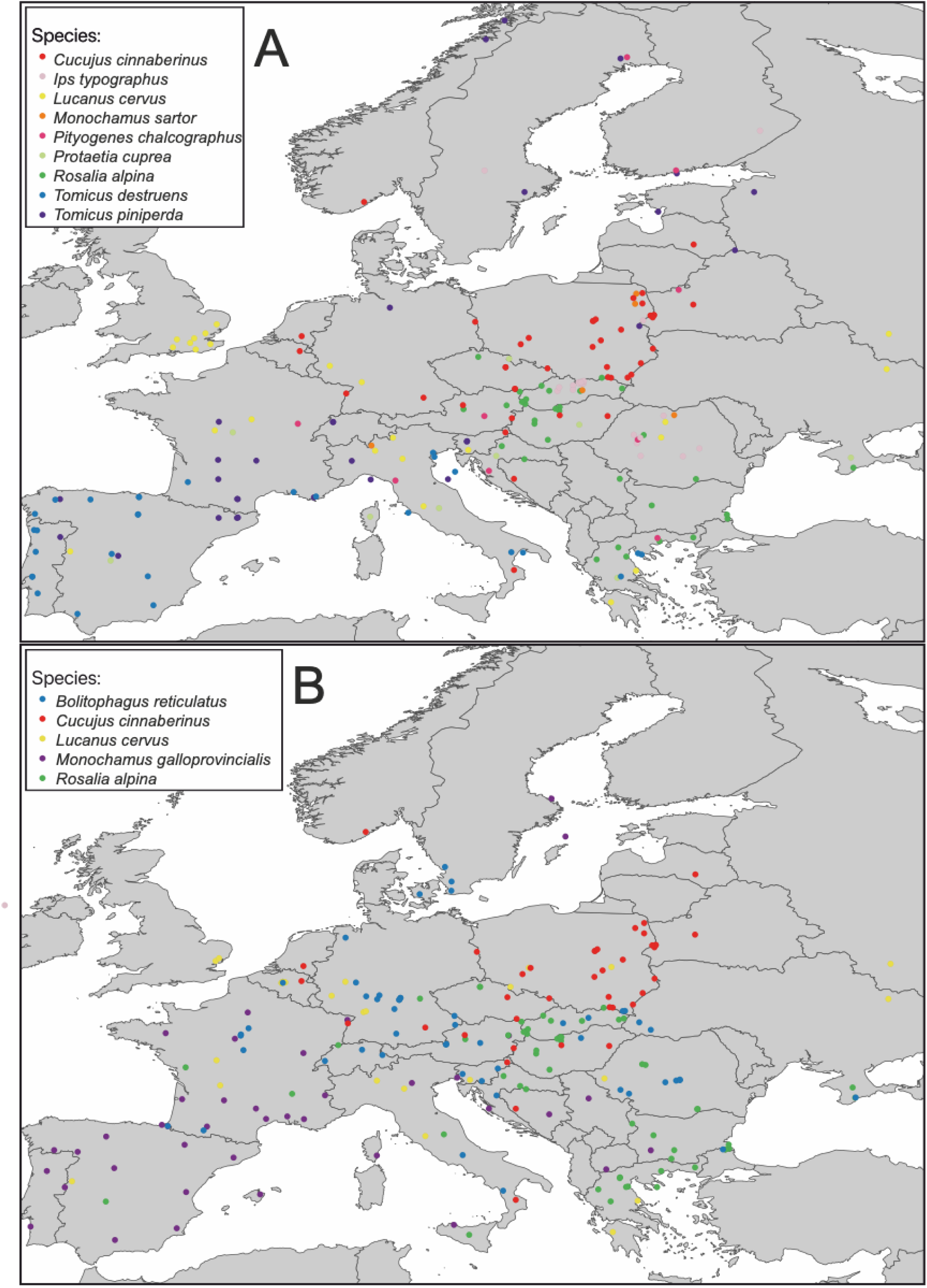
Distribution of sampling sites for nine saproxylic beetle taxa of haplotype diversity (A) and five saproxylic beetle taxa of heterozygosity (B) in Europe

Among the identified articles that met the criteria presented above, only five taxa were available with heterozygosity values from many sites: *Lucanus cervus* (Lucanidae; Solano et al, 2015), *Bolitophagus reticulatus* (Eberle et al. 2021), *Rosalia alpina* (Cerambycidae; Drag et al., 2018), *Monochamus galloprovinciallis* (Cerambycidae; Haran et al., 2018), and *Cucujus cinnaberinus* (Cucujidae; Sikora et al., 2022) (Fig. 1, Table 1, Table S3).

The significant effect of species, latitude, longitude, altitude, and their interactions on genetic diversity (Hd and Ho) were tested. First, model selection analysis (AIC) was done to keep only the most relevant variables (Table 2, Table S4). The results showed a significant effect of the main variables of species, latitudes, and altitudes on haplotype diversity and significant overall main effects of all variables of species, longitudes, latitudes, and altitudes along with some interactions in heterozygosity referring to the leading influence of geographical distribution on genetic diversity with the highest impact on heterozygosity (Table 2, Table S4).

**Table 2.** Effects of species, latitude, longitude, altitude, and their interactions on haplotype diversity (Hd) and heterozygosity (Ho). Significant p-values are in bold. The analyses were limited to the relevant predictors and interactions determined by the model selection analysis.

| Predictor | df | F-value | p-value |
| --- | --- | --- | --- |
| Haplotype diversity data |  |  |  |
| Species | 8 | 6.899 | < 0.001 |
| Latitude | 1 | 14.417 | < 0.001 |
| Altitude | 1 | 8.924 | 0.003 |
| Heterozygosity data |  |  |  |
| Species | 4 | 4.760 | < 0.001 |
| Latitude | 1 | 3.864 | 0.049 |
| Longitude | 1 | 16.312 | < 0.001 |
| Altitude | 1 | 8.601 | 0.003 |
| Latitude:Longitude | 1 | 13.824 | < 0.001 |
| Species: Latitude | 4 | 3.585 | 0.006 |
| Species:Longitude | 4 | 2.340 | 0.052 |

Although geographical distribution showed strong significant effects on genetic diversity, studying the geographical influences of altitude, latitude, and longitude was different between species (Fig. 2, Table 3).

**Fig. 2.**
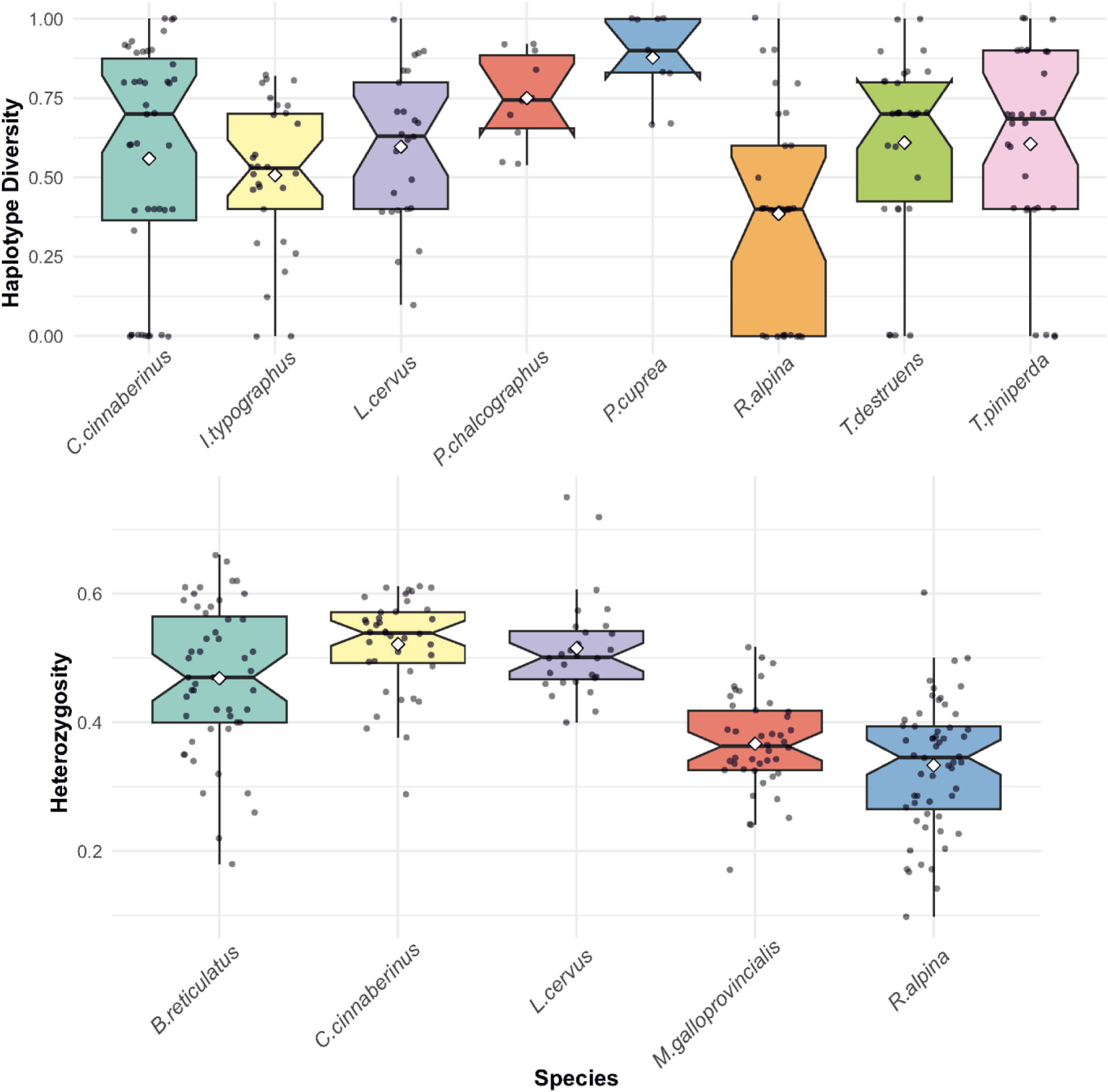
Box-plots visualizing differences in haplotype diversity (A) and heterozygosity (B) of saproxylic beetle taxa in Europe considered for this study

**Table 3.**
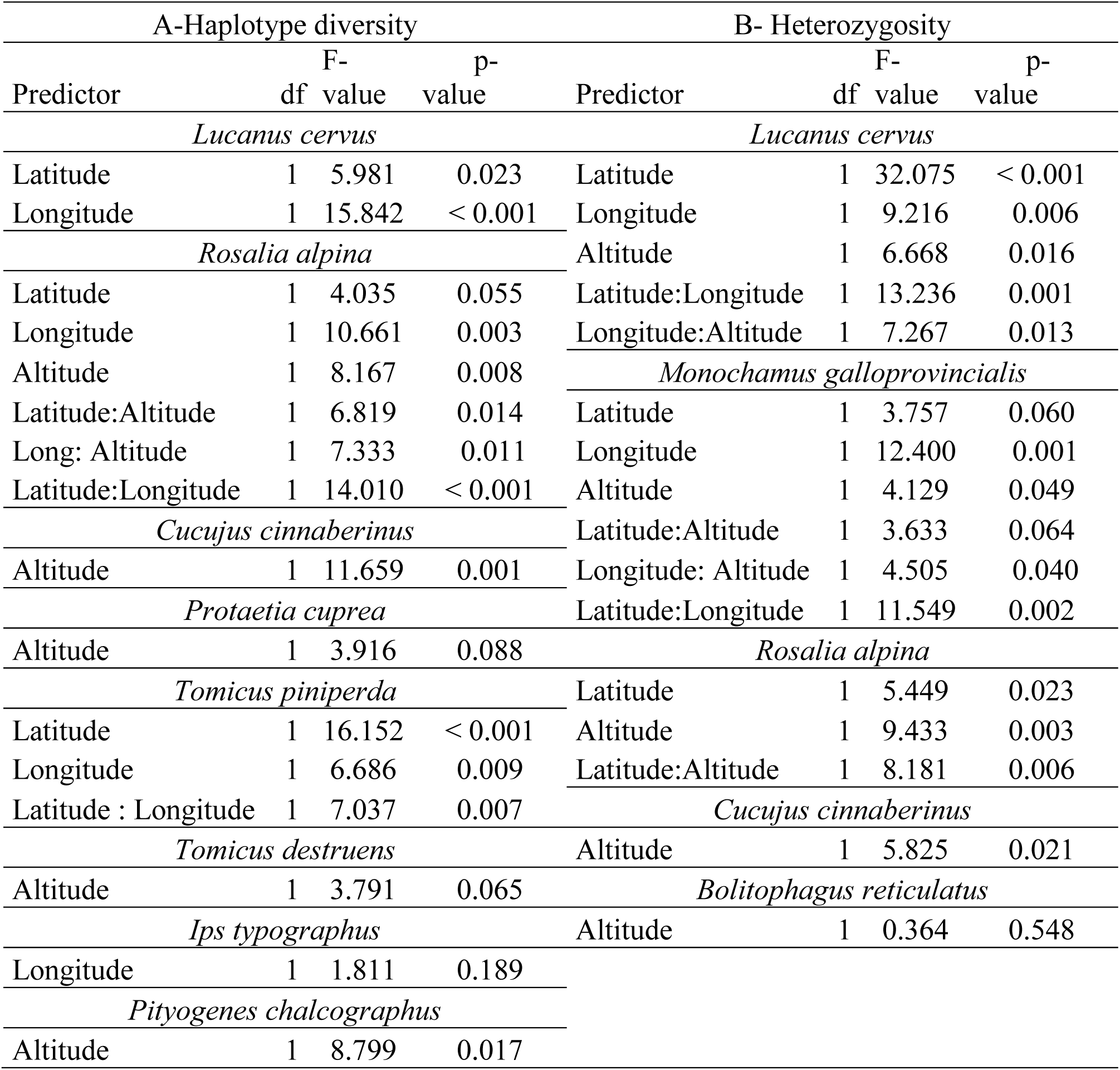
Effects of latitude, longitude, Altitude, and their interactions on A-haplotype diversity and B-heterozygosity of each species separately. The analyses were limited to the relevant predictors and interactions determined by the model selection analysis.

In haplotype diversity, *R. alpina* expressed the lowest diversity values with strong significant effects of geographical distribution of latitude, longitude, and altitude in contrast to *M. sartor* and *P. cuprea* who showed the highest diversity values with the weak significant trend of altitude only in (*P.cuprea*) (Fig. 2, Table 3). By applying model selection analysis to *M. sartor,* the analysis returned “null model”, however, the next best model couldn’t be considered because AIC>2 so, *M. sartor* was excluded from the analysis (Table 3, Table S5).

In heterozygosity, *M. galloprovincialis* and *R. alpina* showed the lowest diversity values with the more significant geographical effects on *M. galloprovincialis* in contrast to *B. reticulatus*, *C. cinnaberinus,* and *L.cervus* who showed the highest diversity values (Fig. 2, Table 3), however, geographical distribution has no significant effect on *B. reticulatus* (Table 3).

The box plots compare haplotype diversity (Hd) and heterozygosity (Ho) across the species considered. The y-axis represents (Hd) or (Ho) respectively, ranging from 0 to 1. Each box shows the interquartile range(IQR), with whiskers extending to 1.5 times the IQR. Individual data points are jittered for clarity, and white diamonds indicate mean values. The notches are used to show confidence intervals around the median.

### 3.3. Correlation with geographic dimensions

The correlation scatter plots revealed varying patterns between species, with some exhibiting strong correlations between latitude and Hd and between altitude and Ho while others showed no discernible trends.

Along latitude following species expressed a decrease of haplotype diversity to the north: *L. cervus*, *R. alpina*, *C. cinnaberinus*, *I. typographus*, *P. chalcographus*, and *T. pinniperda-destruens* (Fig. 3, Fig. S1, S2). In heterozygosity, all species expressed decrease of diversity values to the north except for *B. reticulatus*, however all correlations are insignificant (Fig. 4, Fig. S1, S2)

**Fig. 3.**
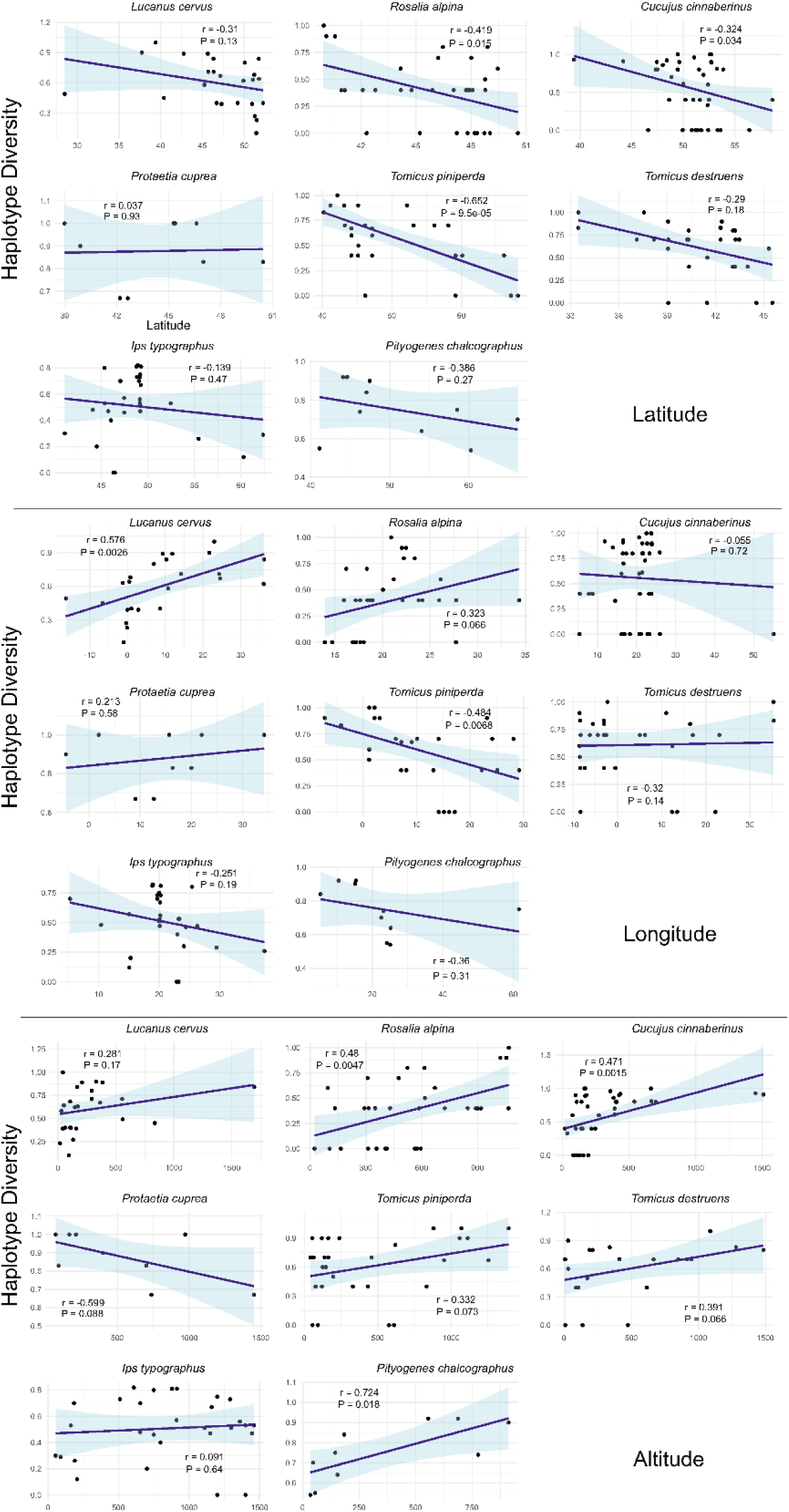
Scatterplots showing changes in haplotype diversity of saproxylic beetles in Europe across latitude (a), longitude (b), and altitude (c). Each plot shows individual data points, a linear regression line with a 95% confidence interval (shaded area), and annotations showing Pearson’s correlation coefficient (r) and p-value.

**Fig. 4.**
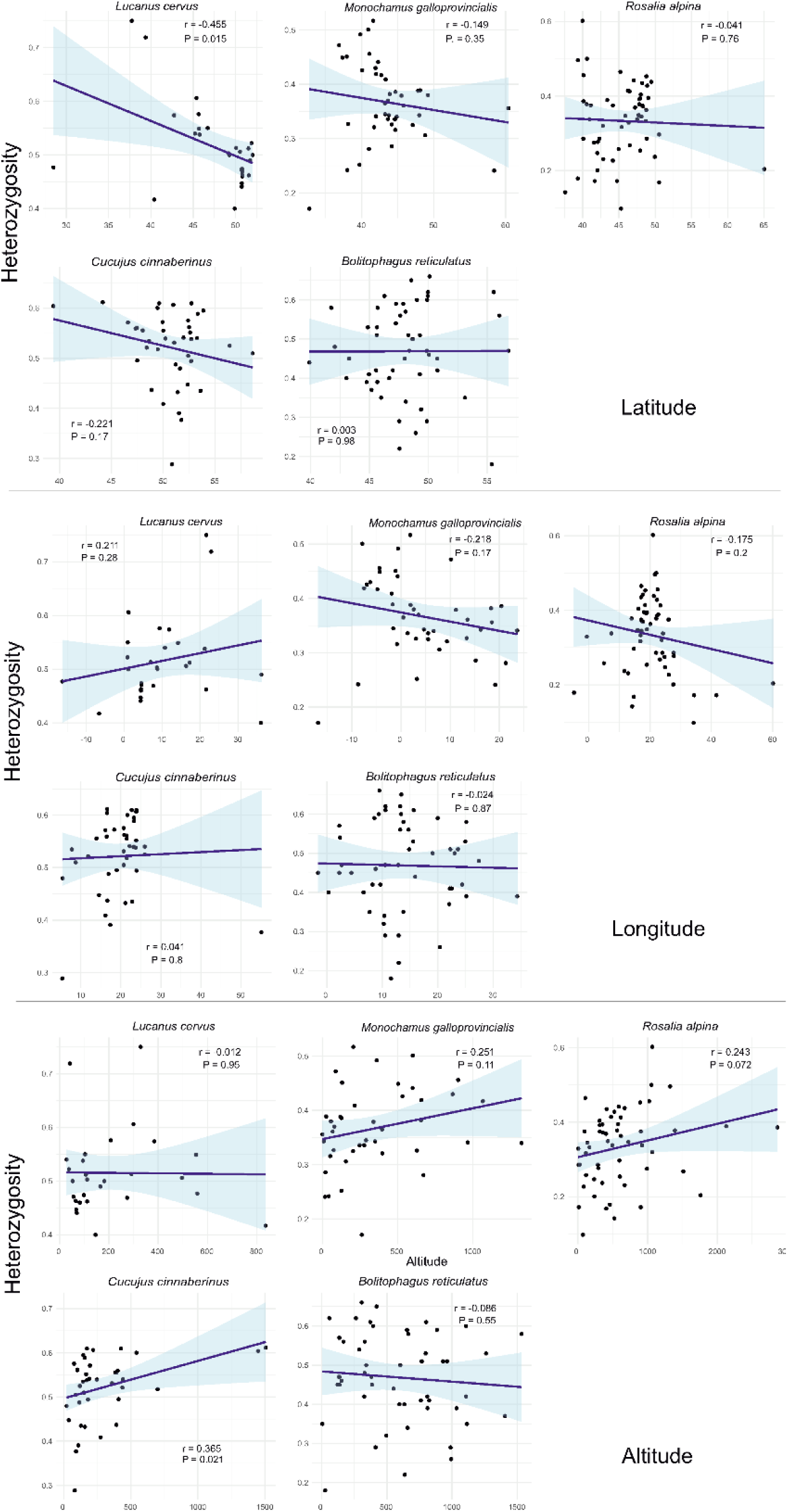
Scatterplots showing changes in heterozygosity of saproxylic beetles in Europe across latitude (a), longitude (b), and altitude (c). Each plot shows individual data points, a linear regression line with a 95% confidence interval (shaded area), and annotations showing Pearson’s correlation coefficient (r) and p-value

Along longitude following species showed decrease in haplotype diversity to the west. For example, *L. cervus*, *R. alpina* and *C. cinnaberinus*. Species that showed increase in haplotype diversity to the west are *I. typographus*, *P. chalcographus*, and *T. pinniperda-destruens* (Fig. 3, Fig. S1, S2). However, for heterozygosity, results revealed either insignificant decrease or increase in diversity values to the west in all species (Fig. 4, Fig. S1, S2)

With increasing altitude only one species showed a decrease in haplotype diversity which is *Protetia cuprea*. *Cucujus cinnaberinus*, *Rosalia alpina*, *Tomicus piniperda* and *Pityogenes chalcographus* showed significant positive correlation with respect to altitude. The rest of the species even though showed positive correlation, the results were not significant (Fig. 3, Fig. S1, S2). For heterozyogisity, only *C. cinnaberinus* showed significant positive correlation with altitude (Fig. 4, Fig. S1, S2).

### 3.4. Genetic units

Information about population structure with the presence of distinct genetic units (clades) was available for 10 species and 7 species complexes

#### 3.4.1. Mitochondrial DNA

The largest number of these taxa have genetic information from following regions: Alpine (18), Balkans (16), Appennines (15), Scandinavia (15), Pannonian-Carpathian-Bohemian (13), Western (12), Eastern (11), North-Western (11), North-Eastern (9), Iberian (9), Anatolian (8), Pontic (6), and Caucasus (3) (Fig. 5a).

**Fig. 5.**
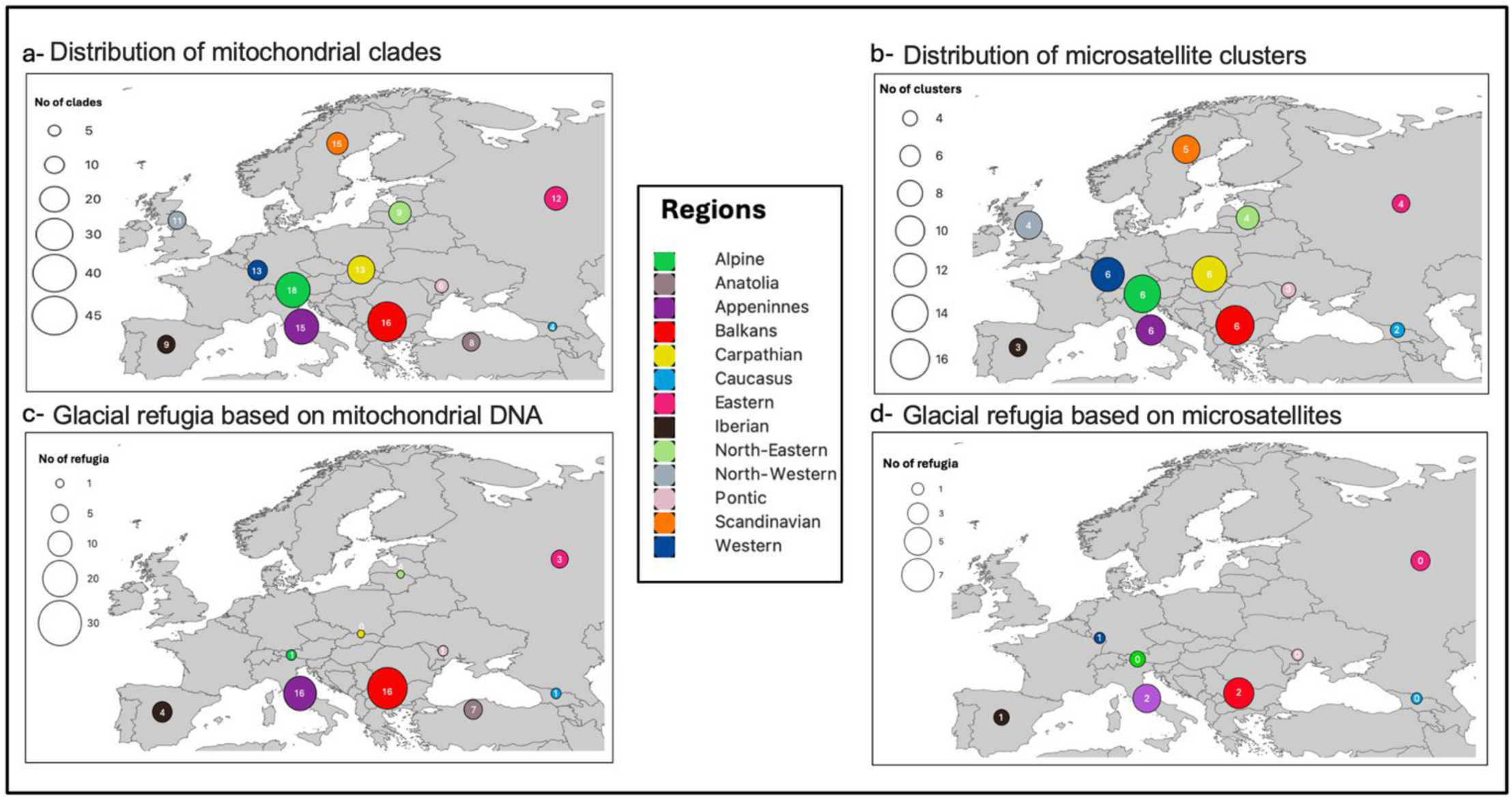
Upper - maps showing the number of mitochondrial clades (a), and microsatellite clusters (b). Numbers in circles represent the number of taxa with data available in particular regions of Europe. Lower - maps showing glacial refugia identified based on mitochondrial DNA (c), and microsatellites (d). Numbers in circles represent the number of unique (private) clades or clusters found in a given region of Europe. Colours indicating different regions of Europe (defined in the method section). Sizes of circles correspond to the numbers of clades (a), clusters (b) or refugia (c,d).

The highest number of evolutionary units was found in the Balkans (42 for 16 species), followed by Apennines (34 for 15 species) and Alpine (34 for 18 species). The lower number of clades was found in Pannonia-Carpathians-Bohemia (25 for 13 species), Scandinavia (24 for 15 species), Western (21 for 13 species), followed by Eastern (17 for 12 species), North-Eastern (16 for 9 species), North-Western (13 for 11 species), and Anatolia (11 for 8 species). The lowest number of clades was found in Iberia (11 for 9 species), Pontic (7 for 6 species), and Caucasus (3 for 3 species) (Fig. 5a).

Regarding unique evolutionary units (present in only one region), the most diverse occurred to be both Balkans (16 for 8 species) and Apennines (16 for 9 species), followed by Anatolia (7 for 5 species), Iberia (7 for 3 species), North-Eastern (4 for 1 species), Eastern (3 for 1 species). Single unique clades were found in Alpine, Pontic, and Caucasus (Fig. 5c).

#### 3.4.2. Microsatellites

Based on the literature and population structure of available data for six species (Table 1):

The largest number of these taxa have genetic information from following regions: Western, Apennines, Balkans, Alpine, and Carpathian (6 in each), followed by Scandinavian (5), North-Western, North-Eastern, Eastern (4 in each), Iberian and Pontic (3 in each), and Caucasus (1). No data was available for Anatolia (Fig. 5b).

The highest number of genetic clusters was found in the Balkans (15 for 6 species), followed by Alpine (14 for 6 species), Pannonia-Carpathians-Bohemia (13 for 6 species), Western (12 for 6 species), North-Western (11 for 4 species), Apennines (10 for 6 species), Scandinavia (9 for 5 species), North-Eastern (7 for 4 species), Eastern (5 for 4 species), Iberian (5 for 3 species), Pontic (4 for 4 species) and Caucasus (3 for 1 species) (Fig. 5b).

Regarding unique clusters (present in only one region), the only cases reported were from Balkans (2 for 2 species), Apennines (2 for 2 species), Iberia (1), and Western (1) (Fig. 5d).

### 3.5. Refugia

A similar pattern was like above when analyzing the localization of glacial refugia reported in articles for saproxylic beetles (Table 1).

Regarding mtDNA, in the Balkans, there are identified 30 such refugia (for 16 species, in some taxa more than one), followed by Apennines (21 for 15 species). In Iberia were found 8 refugia (for 7 species in each), in Anatolia 7 refugia (for 8 species), and in Eastern 6 such areas(for 6 species). In Alpine, Caucasus and Pontic were of 2 refugia (for 2 species in each) were determined, and single ones in Pannonia-Carpathians-Bohemia and North-Eastern, whereas any in Scandinavian, North-Western, and Western (Fig. 5c).

Regarding microsatellites, in the Balkans were identified 7 such refugia (for 6 species, in some taxa more than one), followed by Apennines (6 for 5 species), and Eastern (3 for 3 species). In the Alps and Iberia were identified 2 cases (for 2 species in both), and single in Pontic and Western (Fig. 5d).

## 4. Discussion

Saproxylic beetles inhabiting European forests belong to various taxonomic units and trophic guilds. They have very diverse population structures and dynamics and are known to be either susceptible to forest management or reversely - benefit from forest transformation for wood production (Kozak et al 2021). The common factor for all saproxylic beetles is their association with the wood of dying or dead trees (some of them are responsible for killing of trees, whereas the majority utilize deadwood in various stages of decay) (Grove et al., 2002). Considering all these differences we expect that their genetic diversity is varied. Indeed, this summary of available genetic data for saproxylic European beetles revealed diverse patterns, with surprisingly many congruence mostly caused by a past (Pleistocene and Holocene) history of wood-dwelling organisms.

### 4.1. To the north

Determination of higher genetic diversity for south-European populations of the majority of saproxylic beetles is rather an obvious result considering the history of forest-dwelling species since glacial times (Taberlet et al. 1998; Hewitt 2000; Hagge et al., 2019). This concerns mostly taxa associated with deciduous trees which were forced to retreat to the Mediterranean Basin during the Pleistocene and spread to the north during climate warming in Holocene (Jalut et al, 2009). Not only is the higher genetic diversity observed in southern populations consistent with this pattern, but the presence of distinct clades or clusters and identified refugial areas in Mediterranean peninsulas further supports it. The Balkans have historically served as refugia for many saproxylic beetles, a role that continues to be prominent today (Taberlet et al.,1998). Also Apennines played a substantial role in sheltering populations of wood-dwelling beetles during glacials, although the Alps formed a barrier in the expansion of some of them to the north (Drag et al., 2018). Other areas acting as refugees, but on a lower scale, were Anatolia, Iberia, and Eastern Europe (Varga et al., 2009; Bilgin et al., 2011). This begs the question, could the fewer identified clades and refugia in areas such as Eastern Europe be attributed to limited genetic data available for these regions, or possibly to the extinction of local populations? This discrepancy may also be influenced by historical factors, such as extensive deforestation in regions like Iberia (García-Ruiz et al., 2016). Additionally, the absence of a clear south-north pattern in these areas contrasts with taxa originating from the east, particularly those inhabiting the boreal zone with isolated populations in European mountains (Varga et al., 2008).

### 4.2. East-to-west gradient in forest management

South to north decrease of genetic diversity in saproxylic beetles is not the only geographic pattern that can be observed. Relatively many wood-dwelling Coleoptera express also decreased genetic diversity toward their western populations. It is not the rule, as not all saproxylic beetles have ranges covering eastern Europe - this particularly concerns species associated with temperate deciduous trees having ranges not exceeding Central Europe. Surprisingly many taxa express higher genetic diversity in their central, eastern and south-eastern - European populations, One of the reasons for this pattern could be the presence of beetles belonging to various evolutionary units in this part of Europe as a result of contact between populations having refugia in south-west and soutg-east Europe (phenomenon commonly reported in European biogeography, (Hewitt et al., (2011)). This is likely true for various tenebrionid beetles (Fattorini et al., 2012). Another explanation is that beetle populations in eastern and southeastern Europe are in better condition, being more widespread, abundant, and vital, due to inhabiting higher-quality forests (Karpinski et al., 2021). These forests are either natural or closely resemble natural forests, with a substantial amount of dead wood available (Reif and Walentowski, 2008). Indeed, this part of Europe is known for remnants of primeval forests, such as the Białowieża and Pripyat forests in the lowlands, and the Carpathians and the Balkans in the mountains. Additionally, forest management in eastern and southeastern Europe is generally less intensive than in western, southern, and northern Europe (Angelstam et al., 2011). Although there are forests in eastern and southeastern Europe that are planted and heavily logged, many areas are still covered by semi-natural mixed forests that support rich populations of various species, including saproxylic beetles. While remnants of such semi-natural forests also exist in western and southern Europe, they are mostly confined to small, highly isolated mountainous areas. The fragmentation and isolation of these natural, heterogeneous, and mixed forests likely contribute to the generally lower diversity of saproxylic beetle populations in western and southern Europe. Timber logging in these regions is extensive and most forests in these regions are artificially planted resulting in significant deficiency of dead wood, as it is actively removed from commercial forests (Frank et al., 2003; Horák & Rébl, 2013; Karpinski et al., 2021).

### 4.3. Higher is better ?

The latter observation is supported by findings in this review, which demonstrate that beetle populations sampled at higher altitudes exhibit increased genetic diversity. This increased diversity is likely related to the presence of dead wood, which is typically absent except in areas inaccessible to logging. The most diverse beetle populations have been observed in regions such as the Carpathians, the Balkans, and the Alps (Lassauce et al., 2011). This pattern also contributes to the high diversity of mountain populations in the Mediterranean peninsulas, where refugial populations exist, such as in the Apennines and the Balkans (Karpinski et al., 2021). It is not immediately apparent since populations at higher altitudes must endure more challenging conditions, particularly climatic, than those living in lowlands thus experiencing bottlenecks which typically reduce intrapopulation genetic diversity. Nonetheless most populations of saproxylic beetles exhibit higher genetic diversity in mountainous areas. This highlights the importance of living in heterogeneous forests which are less impacted by management and so have large volumes of deadwood.

### 4.4. Rare vs common

One of the assumptions was that rare, threatened species have overall lower genetic diversity due to the isolation of their populations caused by forest fragmentation and lack of essential microhabitats such as deadwood, in commercial forests (De Groot et al., 2019). The genetic diversity of common species is generally assumed to remain unaffected by forest management. However, a comparison of available data reveals that this is not always the case, and no consistent patterns emerge among saproxylic beetle species (Edelmann et al., 2022).

There are instances where rare species exhibit lower genetic diversity compared to most other taxa. This is particularly evident in species associated with specific tree species and habitats, such as *Lucanus cervus, Cerambyx cerdo*, and *Osmoderma* spp., which depend on old oaks, and *Rosalia alpina*, which is linked to old beeches, especially at the edges of their distribution ranges (Seibold et al., 2015). In contrast, *Cucujus cinnaberinus*, despite also being rare, has high genetic diversity. This species inhabits the bark of various tree species, likely enhancing its dispersal abilities (Vrezec et al., 2017). Common beetles typically exhibit relatively high genetic diversity, which reflects their widespread distribution and numerous populations, primarily in commercial forests such as pine or spruce woods. However, there are exceptions to this pattern. For instance, *Ips typographus* shows relatively low genetic variability within its populations. This species is known for undergoing cyclic outbreaks followed by population collapses, which tend to reduce its overall genetic diversity (Stauffer et al., 2013).

### 4.5. Convergences with other forest-dwellers

The patterns discussed above are also evident in other wood-inhabiting organisms. Among the best-known arboreal animals are woodpeckers (Picidae), whose European members display similar patterns to those of saproxylic beetles which are their primary prey (Fayt et al 2005). Generalist species like the great-spotted woodpecker (*Dendrocopos major*) and green woodpecker (*Picus canus*) exhibit high genetic diversity, but this diversity is unstructured across their European ranges (Myczko et al., 2014). In contrast, distinct evolutionary units in central and east Asia are likely separate species (Perktaş & Quintero, 2013). Similarly, the Iberian woodpecker (*P. sharpei*) was previously considered a subspecies of the green woodpecker (Pons et al., 2011). This unstructured genetic diversity pattern is not observed in saproxylic beetles but may be present in other common, widespread generalist species yet to be examined. Rare specialists associated with deciduous trees, such as the white-backed woodpecker (*Dendrocopos leucotos*) (Pons et al., 2021) and the middle-spotted woodpecker (*Dendrocoptes medius*) (Kamp et al., 2019; Schweizer et al., 2022), have distinct subspecies in their Mediterranean and Middle Eastern ranges, differing from their widespread European mainland counterparts. This pattern is shared with saproxylic beetles like *Rosalia alpina* and *Osmoderma spp*. (Chiari et al., 2013) The three-toed woodpecker (*Picoides tridactylus*) shows the least genetic diversity (Zink et al. 2002), mirroring the genetic patterns of its prey, the bark beetle (*I. typographus)* (Stauffer et al. 1999).

A similar example is found in fungi associated with deadwood, such as the wood decay fungus *Fomitopsis pinicola* or *Gloeoporus taxicola*, which shows similar population structure and genetic diversity across different geographical areas (Högberg et al., 1999; Liu et al., 2021). *Bolitophagus reticulatus* shows similarities in genetic diversification of populations with its host fungi *Fomes fomentarius* (Peintner et al. 2019). These similarities are expected, as it has been previously noted that beetles and fungi exhibit many congruences in their taxonomic, functional, and phylogenetic diversities (Thorn et al., 2018). Likely the same similarities could be expected for other saproxylic organims like moths (Jaworski 2018) or lichens (Rinas et al. 2023), but so far there is deficiency of phylogeographic and population genetic studies on other deadwood-dwelling species.

### 4.6. Implications for conservation or management

In addition to expanding our understanding of historical range changes and contemporary population variability of saproxylic beetles, genetic information about these species as shown here could have significant practical applications. Knowledge of the genetic diversity of “pests” can inform effective management strategies for their populations. This is especially relevant for understanding the causes, mechanisms, and biological consequences of outbreaks, which are cyclic events in the life cycles of economically important species such as bark beetles and certain longhorn or jewel beetles (Stauffer et al., 2013; Haack et al., 2010; Ramasamy et al., 2019). Indeed, ongoing research in this area has shown promising results (https://genomicsofoutbreaks.com/).

Large-scale genomic data can also be used to trace the ancestral populations of invasive beetles and to identify species that are morphologically similar but genetically distinct (Cui et al., 2022). Such research has already been conducted for some saproxylic beetles that are invasive in Europe or for European species that have invaded other regions of the world (Wondafrash et al., 2016). Additionally, genetics could aid in the biological control of pest populations by identifying the microbial, biological or chemical factors that can reduce their numbers in a sustainable way (Elnahal et al., 2022).

The most commonly used designation to study the genetics of rare and threatened taxa is by studying the distribution of distinct evolutionary units of conservation value, such as evolutionary significant units (ESUs) (Casacci et al., 2014) and management units (MUs) (Moritz, 1994). ESUs and MUs are essential for identifying populations that require special protection, particularly those that must be preserved to maintain a substantial part of genetic diversity and potentially important adaptive characteristics vital for the survival of populations (Moritz et al., 1994; Fraser et al., 2001). Additionally, understanding gene flow among populations is important for planning the spatial organization of protected forest sites to sustain connectivity among beetle populations (Balkenhol et al., 2017). Knowing the genetic assignment of individuals to populations is necessary for translocations or reintroductions to avoid outbreeding or inbreeding depression in newly established populations (Maschinski et al., 2013).

Apart from species of economic or conservation value, the majority of saproxylic beetles are natural elements of forest ecosystems, playing crucial roles in carbon and nutrient cycling through their participation in the decay of deadwood (Lieutier et al., 2004). Genetics can also be utilized in this context, for instance, to understand which environmental and spatial factors are responsible for sustaining genetic intra- and inter-population diversity. Tools such as landscape genetics are applicable here, although their use for saproxylic beetles in Europe is still relatively new (Bolliger et al., 2010; Manel et al., 2003).

### 4.7. Prospects and perspectives

The current study serves as a stepping stone for further studies which could potentially employ next-generation sequencing (NGS) technologies like restriction site-associated DNA sequencing or genotyping-by-sequencing (Elshire et al., 2011; Peterson et al., 2012, Narum et al., 2013) to study saproxylic beetles based on large datasets of genomic information (e.g. sets of single nucleotide polymorphisms) and at a larger scale, shedding light on their evolutionary history, adaptation mechanisms, and responses to environmental changes.

Next-generation sequencing (NGS) is already a widely used tool for determining cryptic taxa, including their larvae or pupae, and for delimiting undiscovered evolutionary units (including new taxa) through barcoding (Pante et al. 2015). Additionally, genomic data obtained through NGS can enhance our understanding of phylogenetic relationships among closely related species of saproxylic beetles (Hajibabaei et al., 2007; Lui et al., 2022).

As we have seen, some threatened species are associated with veteran trees (like *Rosalia alpina* with beeches, *Lucanus cervus* and *Osmoderma* spp. with oaks (Drag et al., 2015). So called “pests” are usually specialists of particular tree species (e.g. conifers infested by Scolytinae, Avtzis et al. 2019). Therefore one of the principal future directions should be to incorporate information related to the forest quality and structure into understanding genetic diversity of saproxylic beetles – the topic that has not been addressed so far although knowledge about relations of forest environments with alpha and beta diversity of wood-dwelling beetles is widely studied (Müller,et al. 2015).

Genetic data also show substantial isolation of remote populations particularly in case of rare taxa like *Cucujus cinnaberinus* and *Elater ferrugineus* (Oleksa et al. 2015, Sikora et al. 2023), therefore suitable forest habitat continuity (or forest cover fragmentation) should be considered in future studies on gene flow among populations of saproxylic beetles.

In a similar fashion, *Ips typographus* and other bark beetles, known for its cyclic population dynamics, need also be studied using NGS to examine the genetic diversity across different outbreak and non-outbreak phases (Mykhailenko et al., 2023). By sampling individuals from various outbreak regions and time points, we could elucidate how population fluctuations affect the species’ genetic structure and adaptive potential.

This study opens up various avenues and gives direction for a comprehensive understanding of the genetic dynamics, ecological roles, and conservation needs in the face of ongoing environmental changes of these enigmatic species.

## Author Contributions

The study was designed by Ł. Kajtoch, M. Oczkowicz and R.S.Krovi. Material preparation and data collection were performed by R.S.Krovi. Analyses and visualization were conducted by R.S.Krovi and N. Amer. The first draft of the manuscript was written by R.S.Krovi and Ł.Kajtoch and all authors commented on previous versions of the manuscript. All authors read and approved the final manuscript.

## Data Availability

The datasets generated during and/or analysed during the current study are available in the in Open Science Framework repository under the link: https://osf.io/hsb2c/files/osfstorage/666bfe6865e1de5a73894052

## Supplementary files to

Figure S1. Surface plots showing changes in haplotype diversity and heterozygosity of saproxylic beetles in Europe across latitude, longitude, and altitude.

Files available in Open Science Framework repository under the link: https://osf.io/hsb2c/files/osfstorage/666bfe6865e1de5a73894052

**Figure S2.**
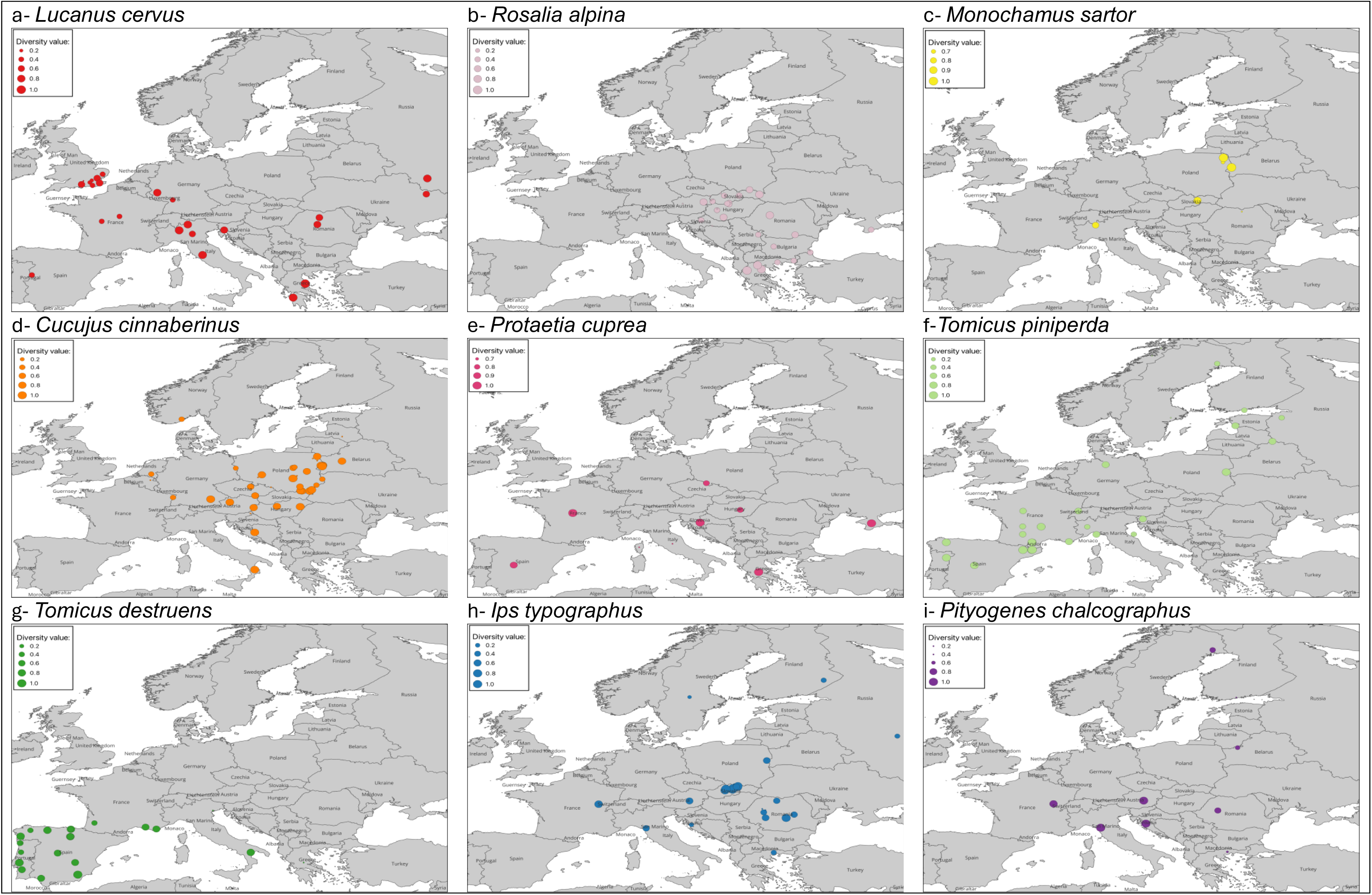
Maps showing the haplotype diversity in nine species of saproxylic beetles collected from different sites in Europe.

**Figure S3.**
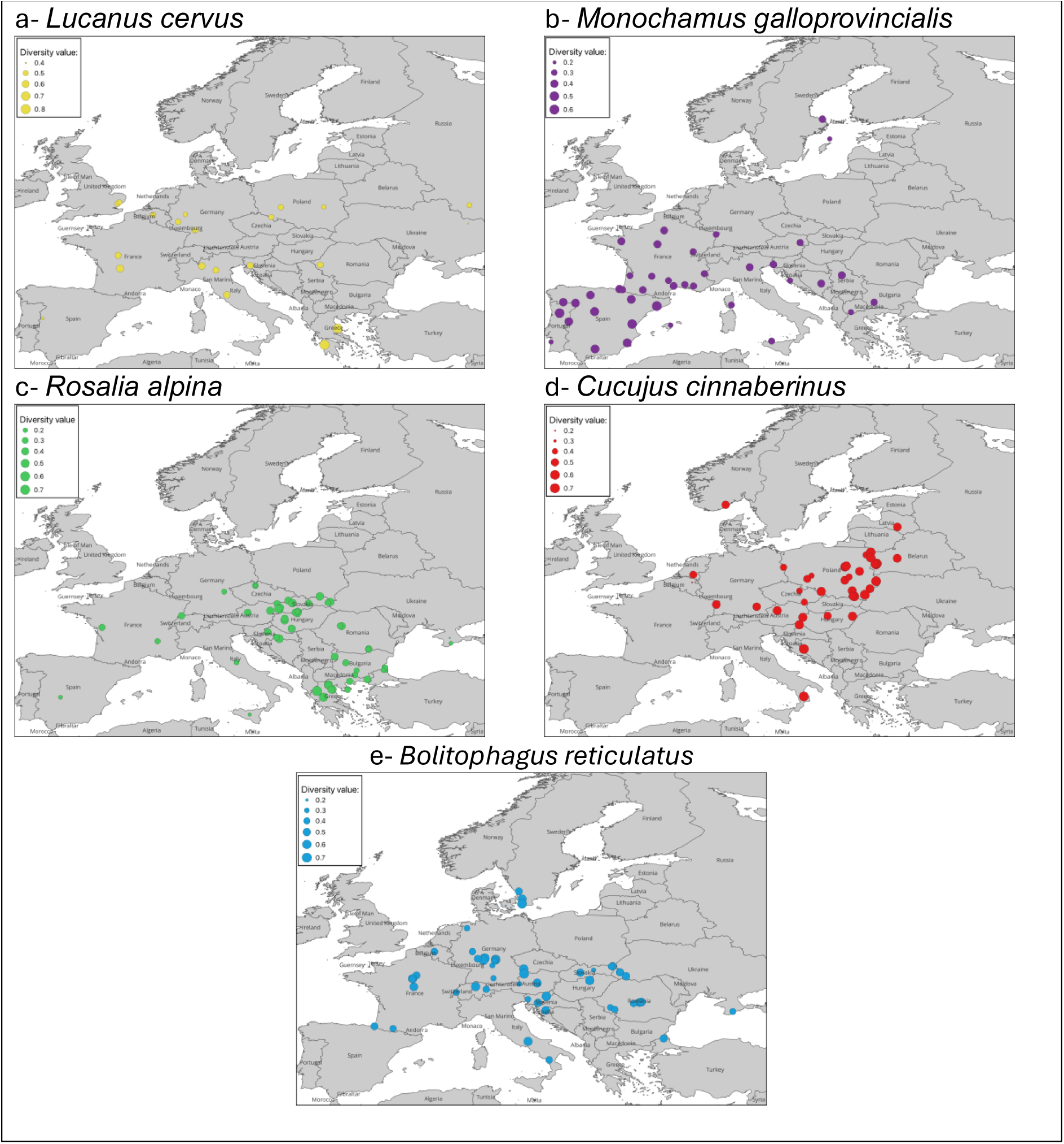
Maps showing heterozygosity in five species of saproxylic beetles collected from different sites in Europe.

**Table S1.**
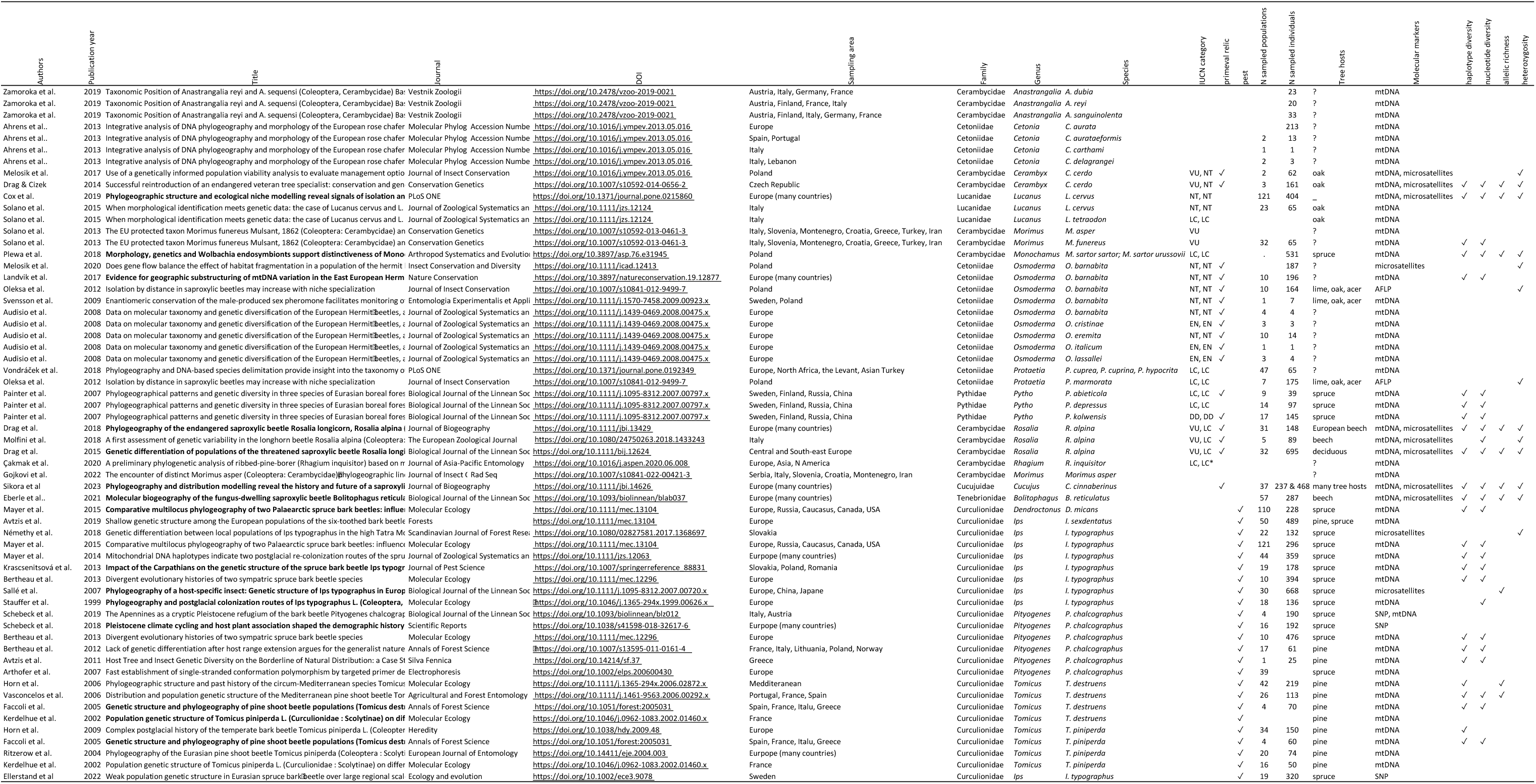
Summary of selected articles retrieved from Web of Science and Scopus databases. Only selected publications referring to saproxylic beetles with phylogeographic or population genetic data are presented. In bold - articles used for meta-analysis.

**Table S2.**
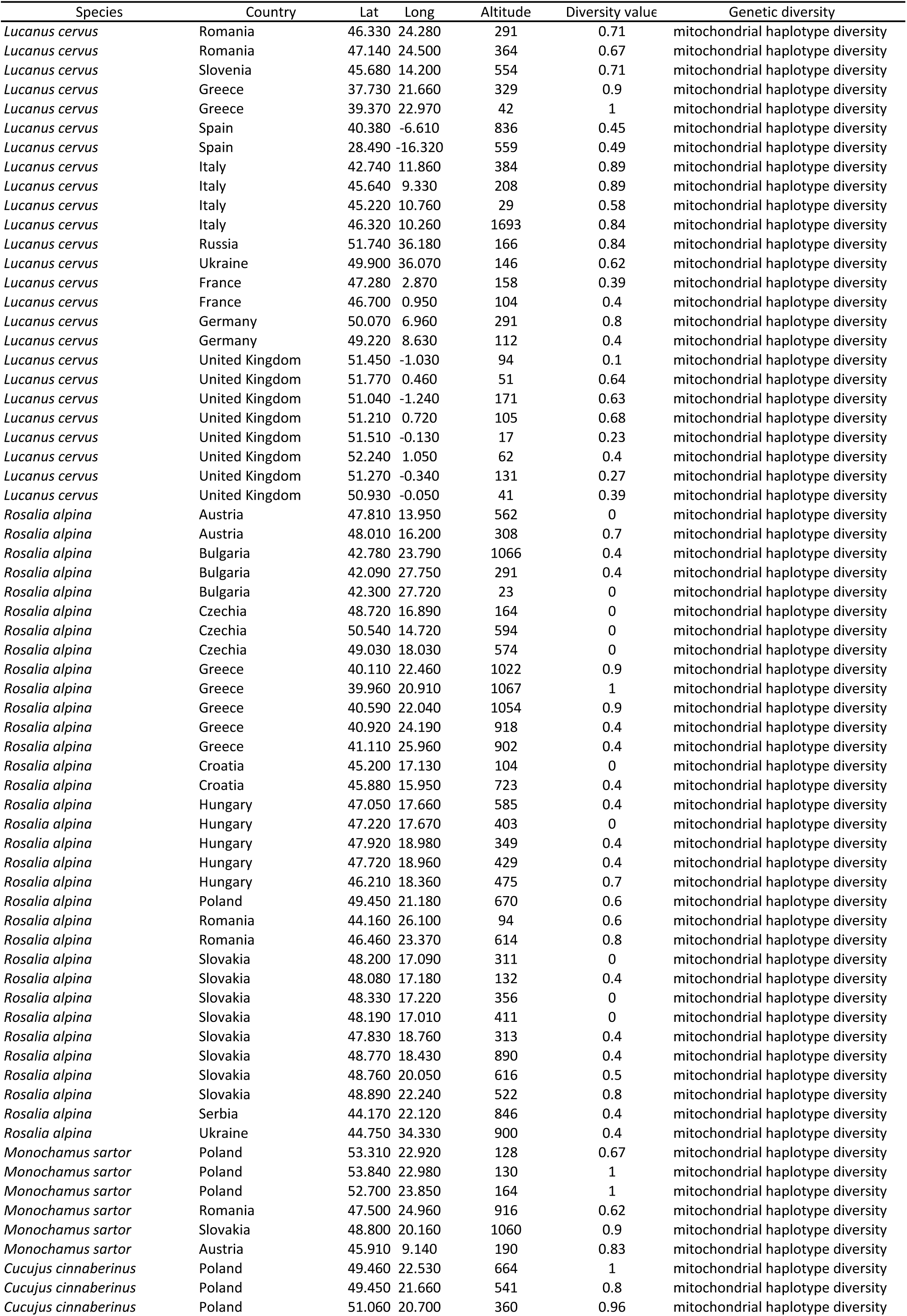
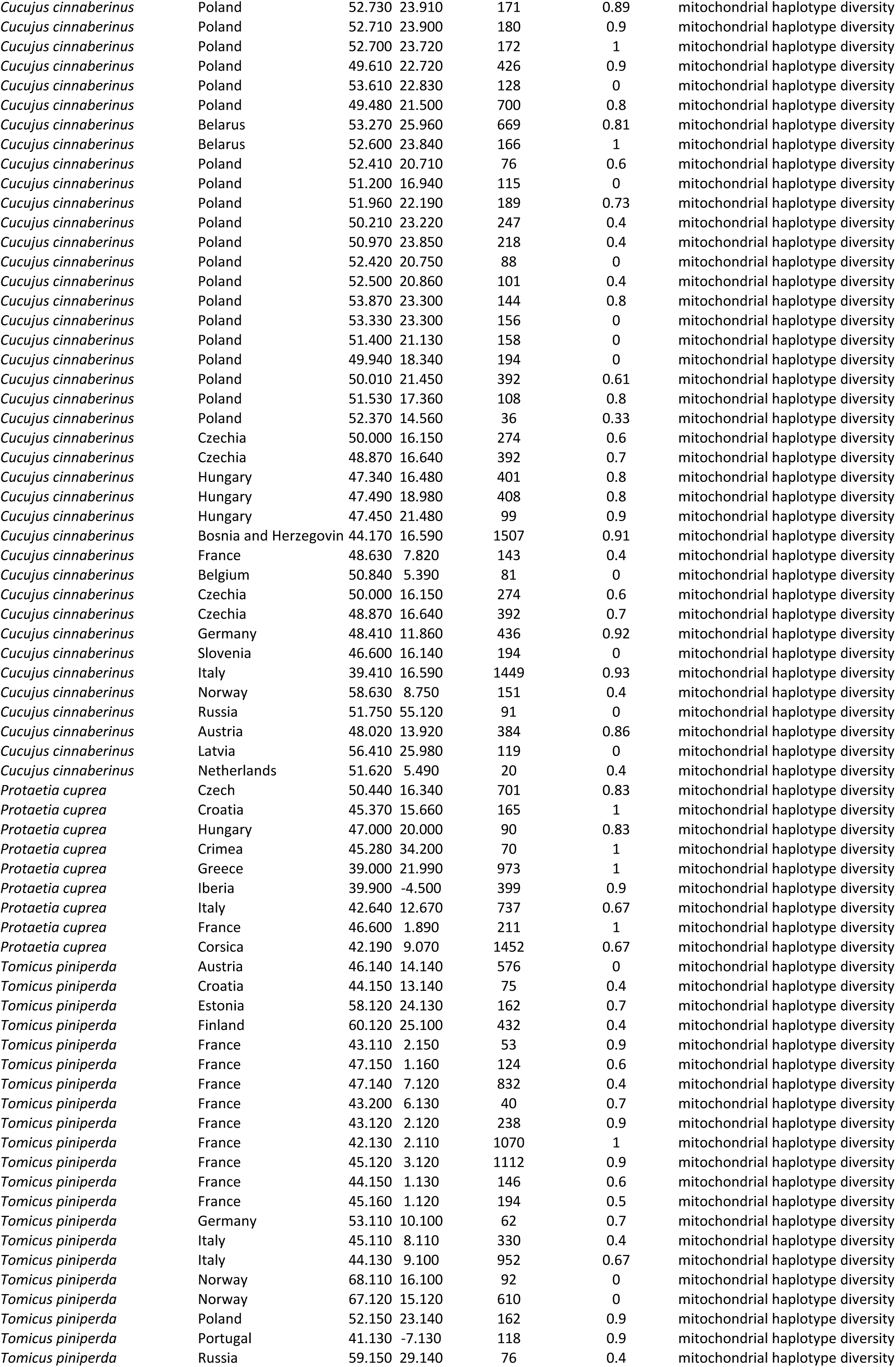
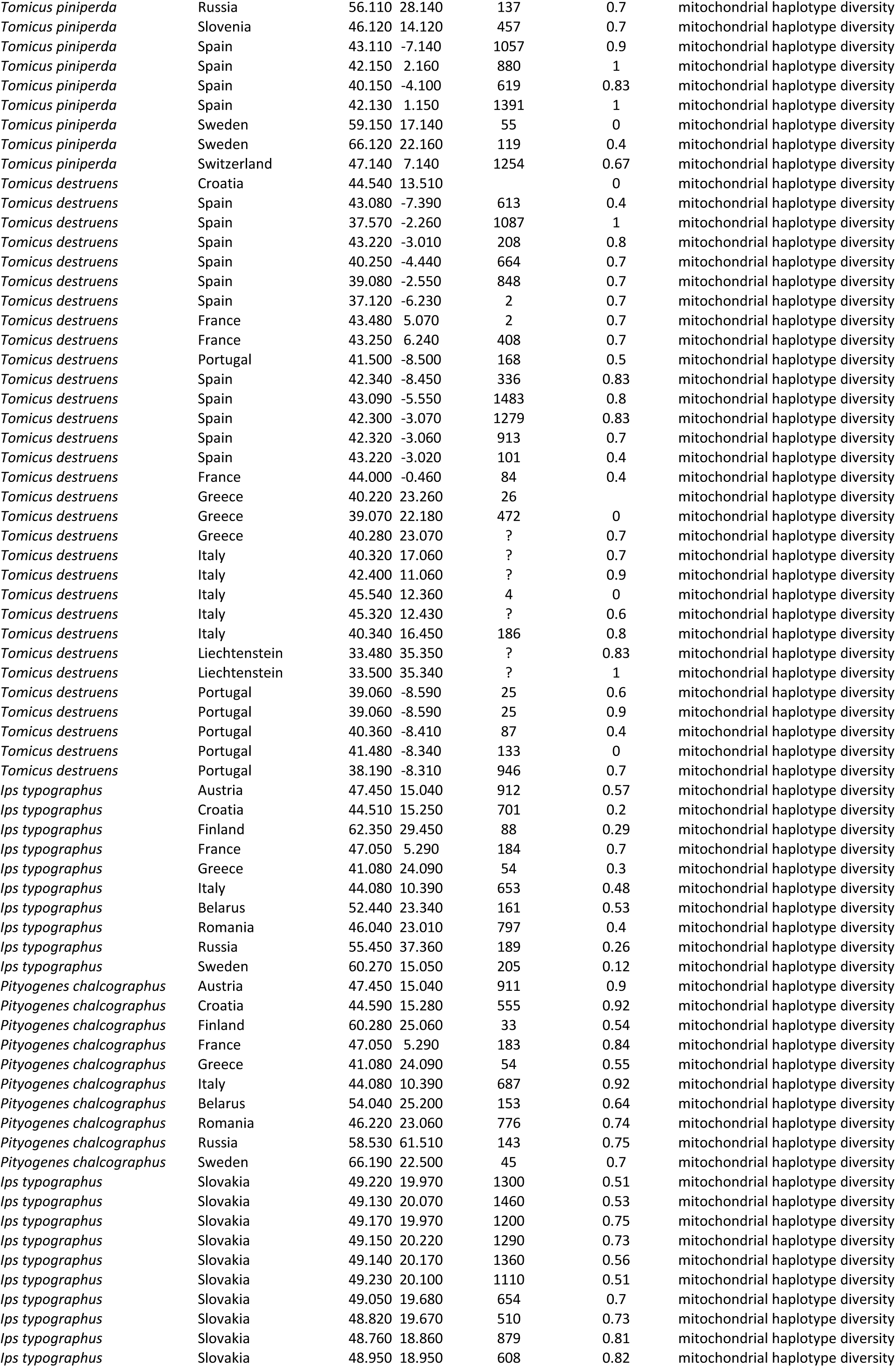
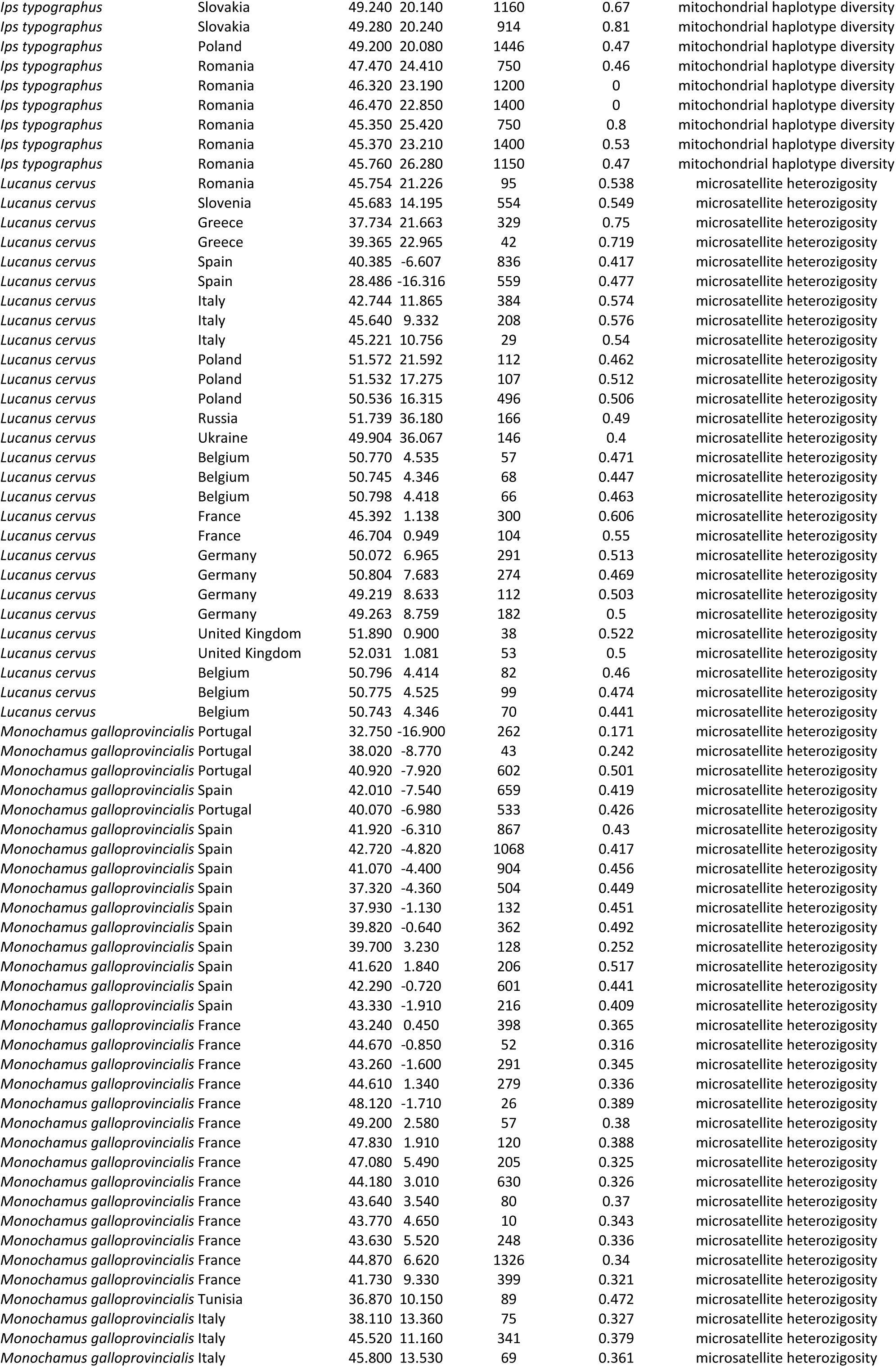
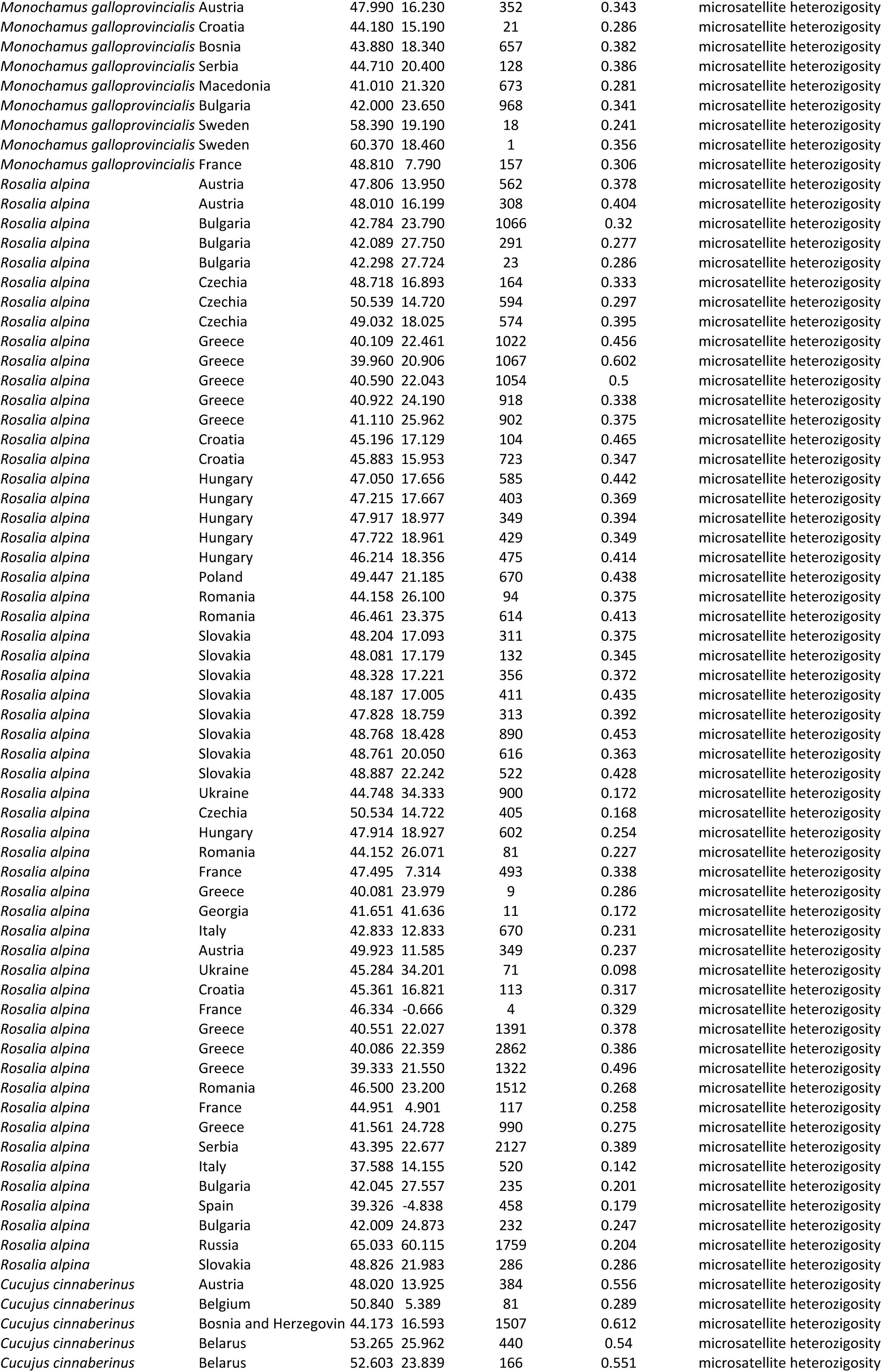
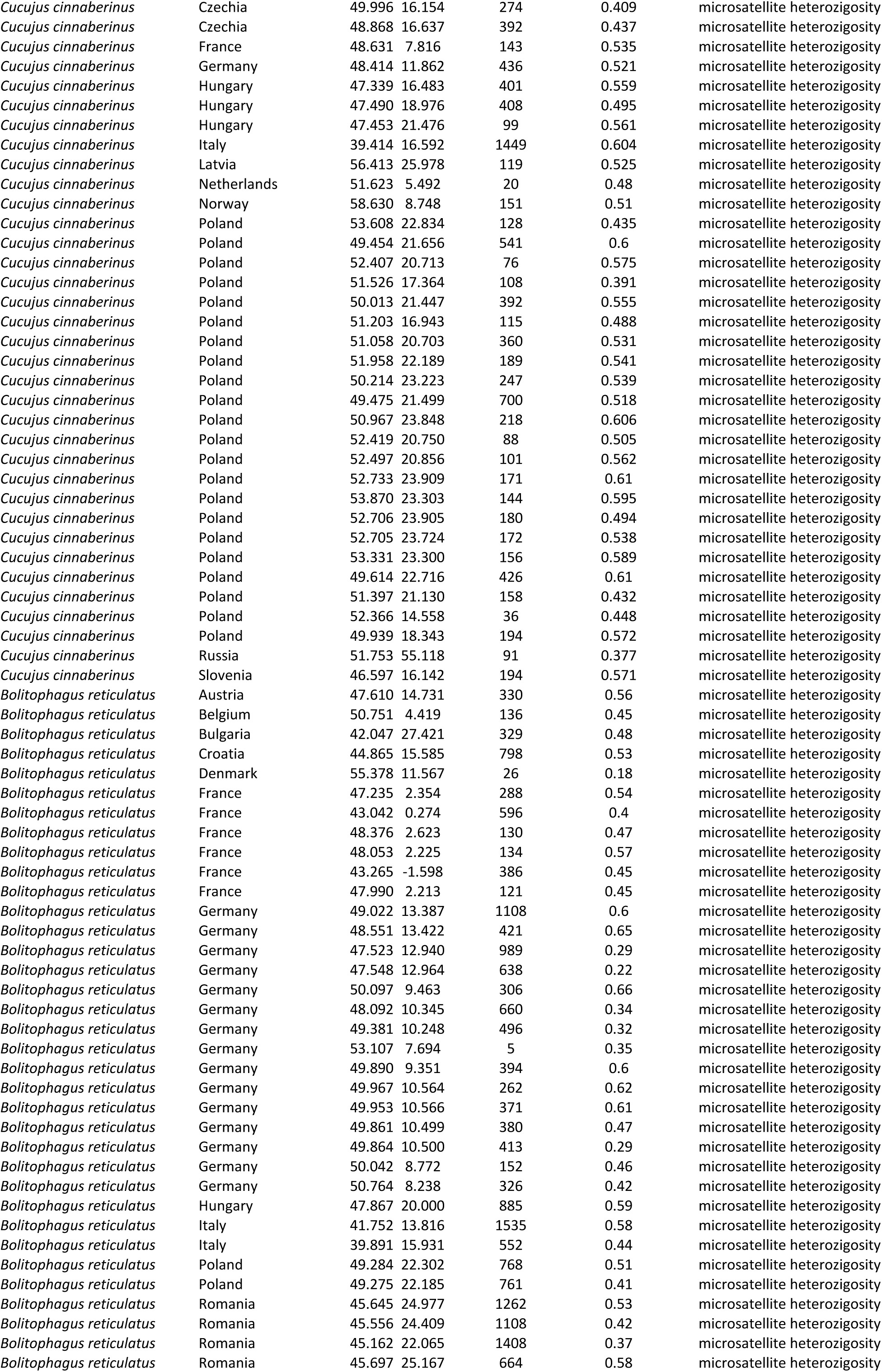
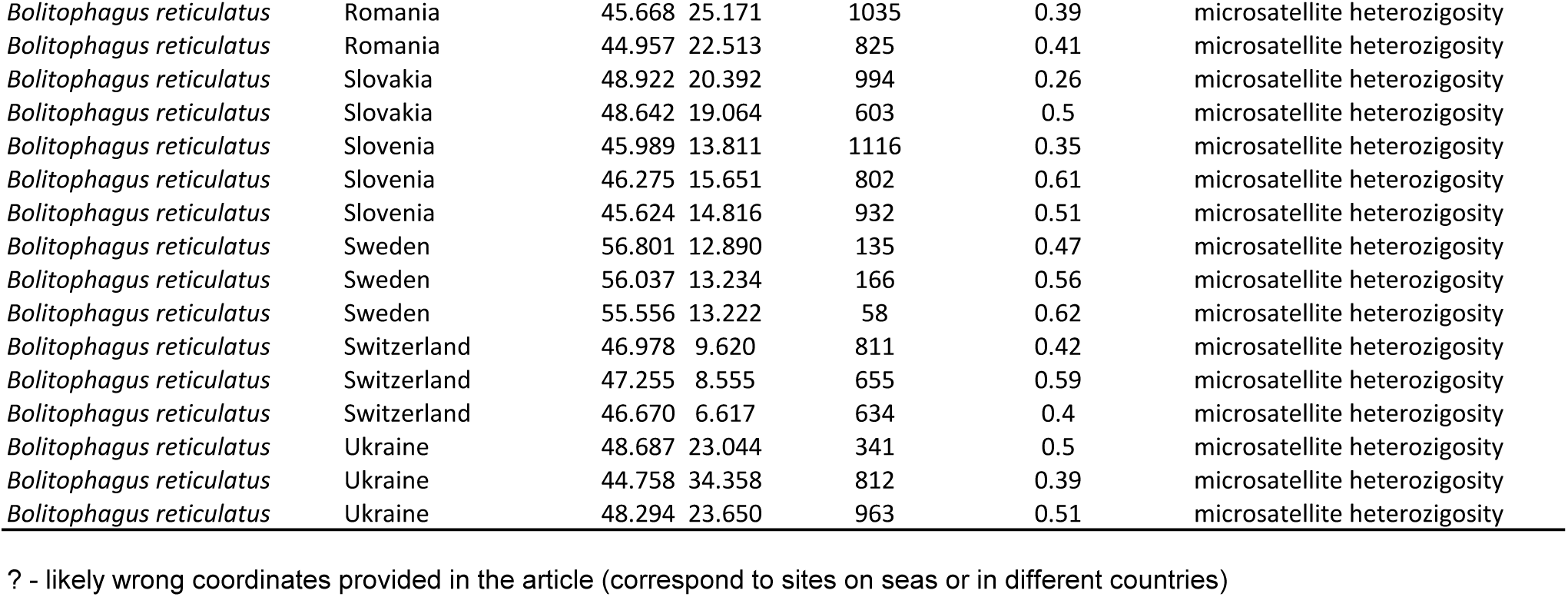
Summary of geographic and genetic data collected from literature about saproxylic beetles from Europe.

**Table S3.** Basic statistics showing mean, maximum, minimum, and standard deviation of Haplotype diversity and heterozygosity.

| Species | mean | min | max | SD |
| --- | --- | --- | --- | --- |
| Haplotype Diversity |  |  |  |  |
| *Cucujus cinnaberinus* | 0.56 | 0.00 | 1.00 | 0.362 |
| *Ips typographus* | 0.51 | 0.00 | 0.82 | 0.236 |
| *Lucanus cervus* | 0.60 | 0.10 | 1.00 | 0.237 |
| *Monochamus sartor* | 0.84 | 0.62 | 1.00 | 0.163 |
| *Pityogenes chalcographus* | 0.75 | 0.54 | 0.92 | 0.144 |
| *Protaetia cuprea* | 0.88 | 0.67 | 1.00 | 0.137 |
| *Rosalia alpina* | 0.39 | 0.00 | 1.00 | 0.308 |
| *Tomicus destruens* | 0.61 | 0.00 | 1.00 | 0.292 |
| *Tomicus piniperda* | 0.61 | 0.00 | 1.00 | 0.312 |
| Heterozygosity |  |  |  |  |
| *Bolitophagus reticulatus* | 0.47 | 0.18 | 0.66 | 0.115 |
| *Cucujus cinnaberinus* | 0.52 | 0.29 | 0.61 | 0.073 |
| *Lucanus cervus* | 0.52 | 0.40 | 0.75 | 0.078 |
| *Monochamus galloprovincia* | 0.37 | 0.17 | 0.52 | 0.074 |
| *Rosalia alpina* | 0.33 | 0.10 | 0.60 | 0.101 |

**Table S4.** Results of the model selection analysis for Diversity values for each haplotype diversity and heterozygous diversity data. Model selection analysis was based on the Akaike’s information criteria for sample size (AICc) and weight as criteria to determine the best explanatory linear model (indicated in red).

| Model | df | logLik | AICc | delta | weight |
| --- | --- | --- | --- | --- | --- |
| Haplotype diversity |  |  |  |  |  |
| **Species+ Latitude+Altitude** | **12** | **-14.04** | **53.67** | **0.00** | **0.52** |
| Species+ Latitude + Altitude +Species:Altitude | 20 | -5.69 | 55.87 | 2.20 | 0.17 |
| Species+Latitude+Altitude+Latitude:Altitude | 13 | -14.03 | 55.94 | 2.27 | 0.17 |
| Species+Latitude+Altitude+Latitude:Altitude+Species:Altitude | 21 | -4.70 | 56.37 | 2.70 | 0.14 |
| Heterozygosity |  |  |  |  |  |
| **Species+Latitude+ Altitude+Longitude+ Species:Latitude+ Species:longitude+ Latitude:Longitude** | **18** | **226.38** | **-413.31** | **0.00** | **0.37** |
| Species+Latitude+Altitude+ Longitude+Species:latitude+Latitude:longitude | 14 | 221.42 | -412.76 | 0.55 | 0.28 |
| Species+Latitude+ Altitude+Longitude+ Species:Latitude+ Species:longitude+ Latitude:Longitude+Altitude | 19 | 226.91 | -411.96 | 1.35 | 0.19 |
| Species+Altitude | 7 | 213.10 | -411.67 | 1.64 | 0.16 |

**Table S5.**
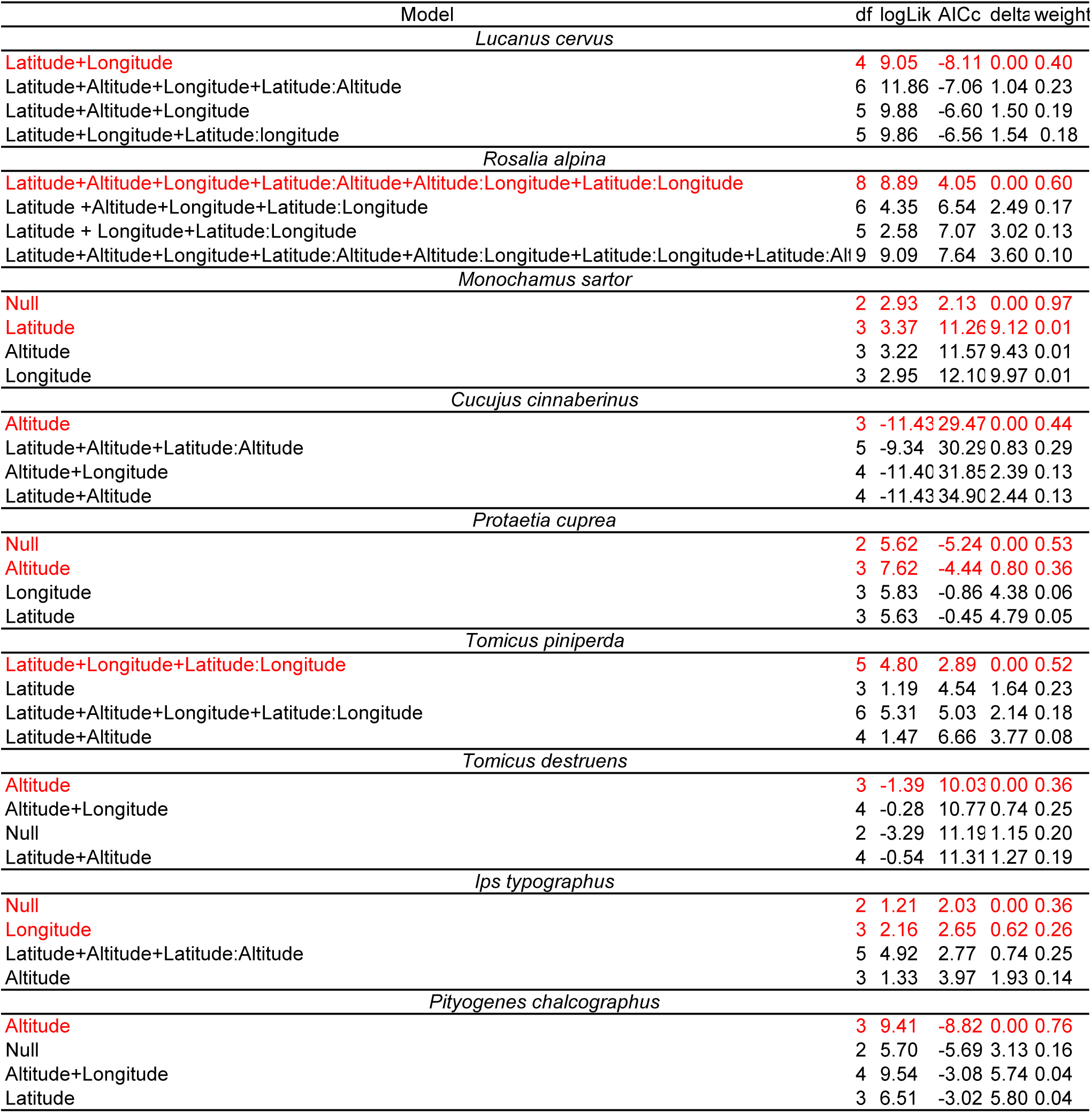
Results of the model selection analysis for haplotype diversity values for each species separately. Model selection analysis was based on the Akaike’s information criteria for sample size (AICc) and weight as criteria to determine the best explanatory linear model (indicated in red).

| Model | df | logLik | AICc | delta | weight |
| --- | --- | --- | --- | --- | --- |
| *Lucanus cervus* |  |  |  |  |  |
| Latitude+Longitude | 4 | 9.05 | -8.11 | 0.00 | 0.40 |
| Latitude+Altitude+Longitude+Latitude:Altitude | 6 | 11.86 | -7.06 | 1.04 | 0.23 |
| Latitude+Altitude+Longitude | 5 | 9.88 | -6.60 | 1.50 | 0.19 |
| Latitude+Longitude+Latitude:longitude | 5 | 9.86 | -6.56 | 1.54 | 0.18 |
| *Rosalia alpina* |  |  |  |  |  |
| Latitude+Altitude+Longitude+Latitude:Altitude+Altitude:Longitude+Latitude:Longitude | 8 | 8.89 | 4.05 | 0.00 | 0.60 |
| Latitude +Altitude+Longitude+Latitude:Longitude | 6 | 4.35 | 6.54 | 2.49 | 0.17 |
| Latitude + Longitude+Latitude:Longitude | 5 | 2.58 | 7.07 | 3.02 | 0.13 |
| Latitude+Altitude+Longitude+Latitude:Altitude+Altitude:Longitude+Latitude:Longitude+Latitude:Alt | 9 | 9.09 | 7.64 | 3.60 | 0.10 |
| *Monochamus sartor* |  |  |  |  |  |
| Null | 2 | 2.93 | 2.13 | 0.00 | 0.97 |
| Latitude | 3 | 3.37 | 11.26 | 9.12 | 0.01 |
| Altitude | 3 | 3.22 | 11.57 | 9.43 | 0.01 |
| Longitude | 3 | 2.95 | 12.10 | 9.97 | 0.01 |
| *Cucujus cinnaberinus* |  |  |  |  |  |
| Altitude | 3 | -11.43 | 29.47 | 0.00 | 0.44 |
| Latitude+Altitude+Latitude:Altitude | 5 | -9.34 | 30.29 | 0.83 | 0.29 |
| Altitude+Longitude | 4 | -11.40 | 31.85 | 2.39 | 0.13 |
| Latitude+Altitude | 4 | -11.43 | 34.90 | 2.44 | 0.13 |
| *Protaetia cuprea* |  |  |  |  |  |
| Null | 2 | 5.62 | -5.24 | 0.00 | 0.53 |
| Altitude | 3 | 7.62 | -4.44 | 0.80 | 0.36 |
| Longitude | 3 | 5.83 | -0.86 | 4.38 | 0.06 |
| Latitude | 3 | 5.63 | -0.45 | 4.79 | 0.05 |
| *Tomicus piniperda* |  |  |  |  |  |
| Latitude+Longitude+Latitude:Longitude | 5 | 4.80 | 2.89 | 0.00 | 0.52 |
| Latitude | 3 | 1.19 | 4.54 | 1.64 | 0.23 |
| Latitude+Altitude+Longitude+Latitude:Longitude | 6 | 5.31 | 5.03 | 2.14 | 0.18 |
| Latitude+Altitude | 4 | 1.47 | 6.66 | 3.77 | 0.08 |
| *Tomicus destruens* |  |  |  |  |  |
| Altitude | 3 | -1.39 | 10.03 | 0.00 | 0.36 |
| Altitude+Longitude | 4 | -0.28 | 10.77 | 0.74 | 0.25 |
| Null | 2 | -3.29 | 11.19 | 1.15 | 0.20 |
| Latitude+Altitude | 4 | -0.54 | 11.31 | 1.27 | 0.19 |
| *Ips typographus* |  |  |  |  |  |
| Null | 2 | 1.21 | 2.03 | 0.00 | 0.36 |
| Longitude | 3 | 2.16 | 2.65 | 0.62 | 0.26 |
| Latitude+Altitude+Latitude:Altitude | 5 | 4.92 | 2.77 | 0.74 | 0.25 |
| Altitude | 3 | 1.33 | 3.97 | 1.93 | 0.14 |
| *Pityogenes chalcographus* |  |  |  |  |  |
| Altitude | 3 | 9.41 | -8.82 | 0.00 | 0.76 |
| Null | 2 | 5.70 | -5.69 | 3.13 | 0.16 |
| Altitude+Longitude | 4 | 9.54 | -3.08 | 5.74 | 0.04 |
| Latitude | 3 | 6.51 | -3.02 | 5.80 | 0.04 |

**Table S6.** Results of the model selection analysis for heterozygous diversity values for each species separately. Model selection analysis was based on the Akaike’s information criteria for sample size (AICc) and weight as criteria to determine the best explanatory linear model.

| Model | df | logLik | AICc | delta | weight |
| --- | --- | --- | --- | --- | --- |
| *Lucanus cervus* |  |  |  |  |  |
| Latitude+Altitude+Longitude+Latitude:Longitude+Altitude:Longitude | 7 | 55.25 | -90.91 | 0.00 | 0.52 |
| Latitude+Altitude+Longitude+Latitude:Altitude+ Latitude:Longitude+Altitude:Longitude | 8 | 56.43 | -89.28 | 1.63 | 0.23 |
| Latitude+Longitude+Latitude:Longitude | 5 | 50.51 | -88.28 | 2.63 | 0.14 |
| Latitude+Altitude+Longitude+Latitude:Altitude+Latitude:Longitude | 7 | 53.70 | -87.80 | 3.11 | 0.11 |
| *Monochamus galloprovincialis* |  |  |  |  |  |
| Latitude+Altitude+Longitude+Latitude:Altitude+Altitude:Longitude+Latitude:Longitude | 8 | 60.18 | -99.99 | 0.00 | 0.45 |
| Latitude +Altitude+Longitude+Altitude:Longitude+Latitude:Longitude | 7 | 58.10 | -98.91 | 1.08 | 0.26 |
| Latitude +Altitude+Longitude+Altitude:Longitude+Latitude:Longitude | 7 | 57.64 | -97.98 | 2.02 | 0.17 |
| Altitude+Latitude+Longitude+Altitude:Latitude+Latitude:Altitude+Latitude:Longitude | 9 | 60.45 | -97.27 | 2.72 | 0.12 |
| *Rosalia alpina* |  |  |  |  |  |
| Latitude+Altitude+Latitude:Altitude | 5 | 55.16 | -99.13 | 0.00 | 0.53 |
| Altitude+Longitude | 4 | 52.89 | -97.00 | 2.12 | 0.18 |
| Latitude+Altitude+Longitude+Latitude:Altitude | 6 | 55.27 | -96.83 | 2.30 | 0.17 |
| Latitude+Altitude+Longitude+Latitude:Longitude | 6 | 54.95 | -96.19 | 2.94 | 0.12 |
| *Cucujus cinnaberinus* |  |  |  |  |  |
| Altitude | 3 | 51.23 | -95.78 | 0.00 | 0.55 |
| Altitude+Longitude | 4 | 51.34 | -93.53 | 2.25 | 0.18 |
| *Bolitophagus reticulatus* |  |  |  |  |  |
| Null | 2 | 38.58 | -72.92 | 0.00 | 0.49 |
| Altitude | 3 | 38.77 | -71.04 | 1.88 | 0.19 |
| Longitude | 3 | 38.60 | -70.69 | 2.23 | 0.16 |
| Latitude | 3 | 38.58 | -70.66 | 2.26 | 0.16 |

